## Supplementary Information for "Spatial connectivity increases ecosystem resilience towards an ongoing regime shift"

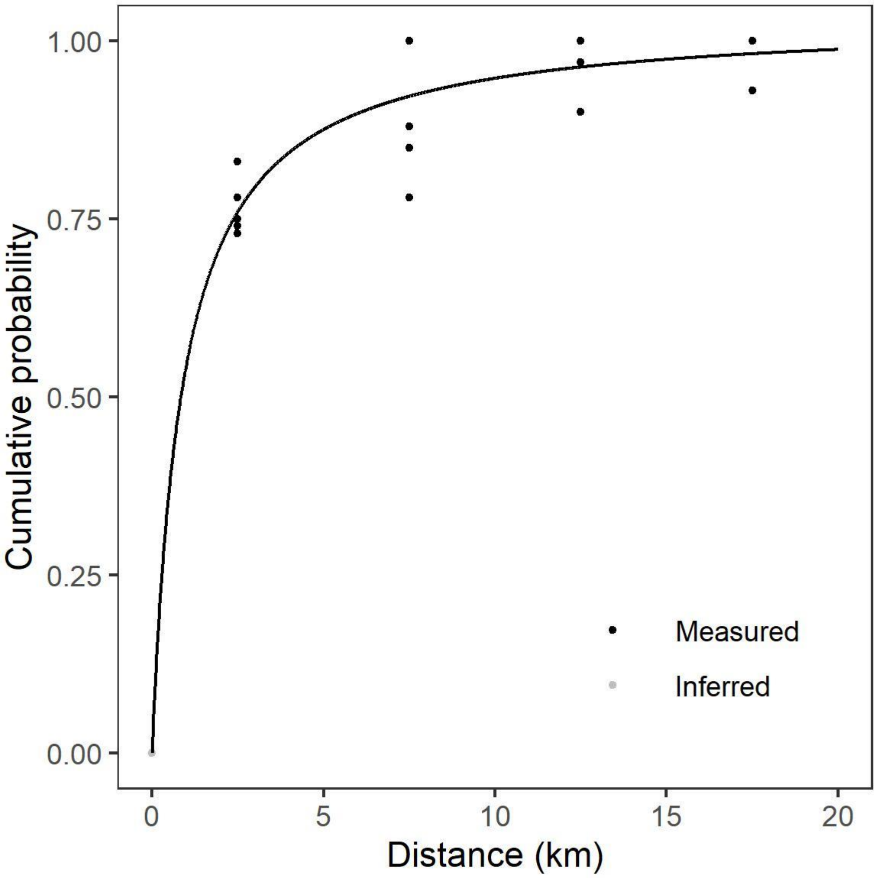


**Supplementary Fig. 1. Dispersal distances for perch.** Points indicate the cumulative proportion of fish caught at a given distance based on tagging data from perch tagged and re-caught in May for several locations in Finland (Siddika & Lehtonen, 2004). The grey points illustrate the straightforward assumption that all dispersal distances were 0 or longer. The black line show the Michaelis-Menten model fit to the data (both grey and black points), showing the predicted cumulative probability of dispersing a given distance, or shorter.

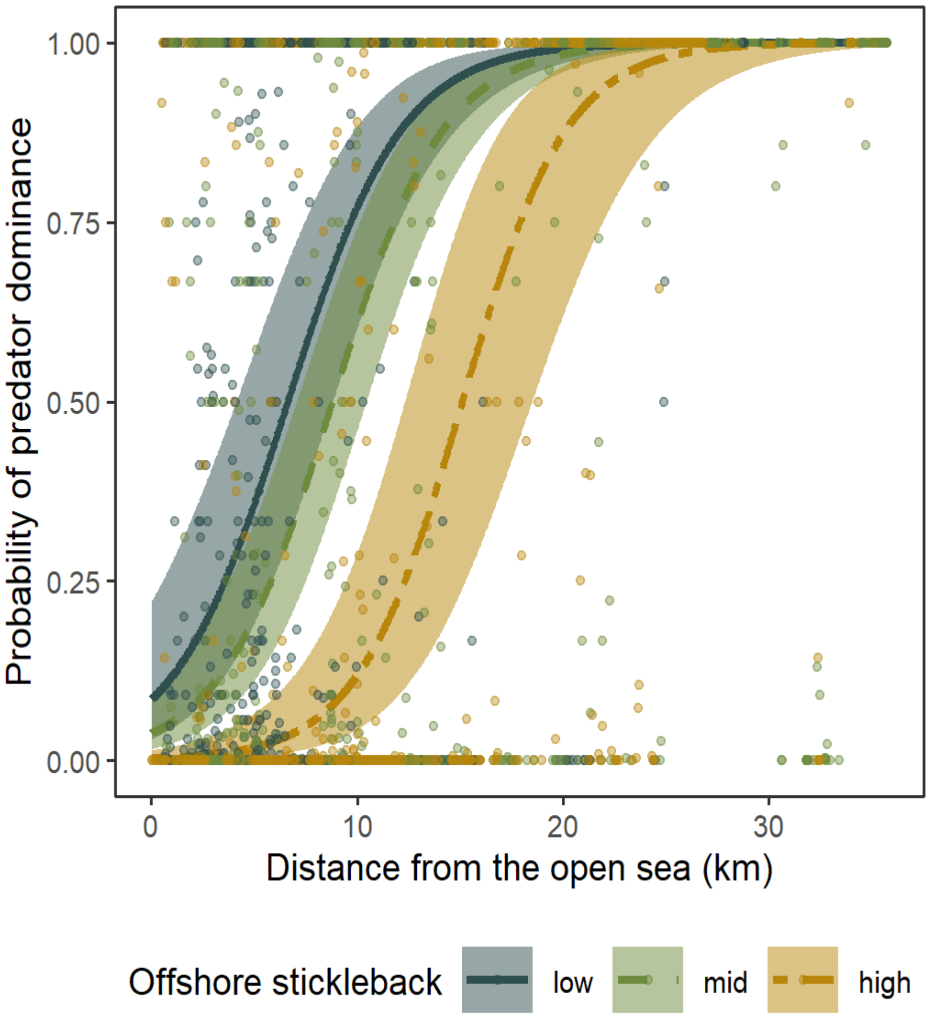


**Supplementary Fig. 2. Probability of predator dominance as a function of distance from the open sea for different offshore densities of mature stickleback.** low = 10th percentile, mid = median, high = 90th percentile. Lines show predictions from a generalised linear mixed model including spatio-temporal random fields and an effect of $\log_{10}$-transformed wave exposure (here at median value), with 95% confidence intervals. Points show raw data colour-coded according to offshore stickleback densities (split into 3 equally sized groups for visualisation purposes).

**Supplementary Table 1. Estimated coefficients with 95% confidence intervals from models of predatory fish dominance as a function of incoming stickleback.** Main effects of open sea stickleback densities, distance from the open sea, $log_{10}$-transformed wave exposure, and an interaction between open sea stickleback densities and distance from the open sea, with different random effect structures. All models are fitted to the same dataset (N = 3491). Estimates are based on scaled variables. See Supplementary Fig. 2 for a visualisation of the effects of distance to the open sea and open sea stickleback densities. Values are coloured in green if lower and upper limit both indicate a positive effect, and red if lower and upper limit both indicate a negative effect.

| Model | Effect | | | |
|---|---|---|---|---|
| | distance to open sea | open sea stickleback | distance × stickleback | $log_{10}$(wave exposure) |
| Base model | 1.13 (1.02; 1.24) | -0.43 (-0.51; -0.34) | 0.19 (0.08; 0.30) | -0.15 (-0.23; -0.07) |
| Across-area random effect of year | 1.24 (1.12; 1.37) | -0.80 (-0.93; -0.67) | 0.27 (0.14; 0.39) | -0.29 (-0.38; -0.20) |
| Constant spatial random fields | 2.66 (1.94; 3.37) | -0.44 (-0.60; -0.28) | -0.13 (-0.34; 0.08) | -0.53 (-0.73; -0.33) |
| Constant spatial random fields and across-area random effect of year | 2.88 (2.07; 3.68) | -0.36 (-0.62; -0.09) | 0.08 (-0.16; 0.32) | -0.68 (-0.90; -0.46) |
| Year-specific spatial random fields | 3.23 (2.64; 3.81) | -1.20 (-1.77; -0.63) | 0.11 (-0.38; 0.59) | -0.73 (-0.95; -0.51) |

**Supplementary Table2. Estimated coefficients with 95% confidence intervals from models of predatory fish densities as a function of incoming stickleback.** Main effects of open sea stickleback densities, distance from the open sea, $log_{10}$-transformed wave exposure, and an interaction between open sea stickleback densities and distance from the open sea, with different random effect structures. All models are fitted to the same dataset (N = 7415). Estimates are based on scaled variables. Values are coloured in green if lower and upper limit both indicate a positive effect, and red if lower and upper limit both indicate a negative effect.

| Model | Effect | | | |
|---|---|---|---|---|
| | distance to open sea | open sea stickleback | distance × stickleback | $log_{10}$(wave exposure) |
| Base model | 0.74 (0.65; 0.83) | -0.21 (-0.30; -0.12) | 0.19 (0.08; 0.29) | 0.01 (-0.08; 0.10) |
| Across-area random effect of year | 0.78 (0.69; 0.88) | -0.15 (-0.26; -0.03) | 0.31 (0.20; 0.42) | -0.28 (-0.39; -0.17) |
| Constant spatial random fields | 1.26 (0.84; 1.68) | -0.09 (-0.21; 0.02) | 0.04 (-0.08; 0.16) | -0.19 (-0.36; -0.02) |
| Constant spatial random fields and across-area random effect of year | 1.24 (0.79; 1.69) | 0.15 (0.00; 0.31) | 0.25 (0.10; 0.39) | -0.30 (-0.47; -0.13) |
| Year-specific spatial random fields | 1.24 (0.95; 1.53) | -0.54 (-0.88; -0.20) | 0.16 (-0.10; 0.42) | -0.28 (-0.44; -0.13) |

**Supplementary Table 3. Estimated coefficients with 95% confidence intervals from models of stickleback densities as a function of incoming stickleback.** Main effects of open sea stickleback densities, distance from the open sea, $log_{10}$-transformed wave exposure, and an interaction between open sea stickleback densities and distance from the open sea, with different random effect structures. All models are fitted to the same dataset (N = 7167). Estimates are based on scaled variables. Values are coloured in green if lower and upper limit both indicate a positive effect, and red if lower and upper limit both indicate a negative effect.

| Model | Effect | | | |
|---|---|---|---|---|
| | distance to open sea | open sea stickleback | distance × stickleback | $log_{10}$(wave exposure) |
| Base model | -0.66 (-0.76; -0.56) | 0.61 (0.48; 0.74) | 0.30 (0.17; 0.43) | 0.38 (0.25; 0.50) |
| Across-area random effect of year | -1.26 (-1.41; -1.10) | 0.89 (0.72; 1.06) | 0.35 (0.20; 0.50) | 0.38 (0.24; 0.53) |
| Constant spatial random fields | -2.14 (-2.64; -1.64) | 0.59 (0.41; 0.77) | 0.03 (-0.17; 0.24) | 0.41 (0.20; 0.62) |
| Constant spatial random fields and across-area random effect of year | -2.06 (-2.61; -1.52) | 0.58 (0.32; 0.84) | 0.05 (-0.18; 0.27) | 0.43 (0.22; 0.64) |
| Year-specific spatial random fields | -2.62 (-3.11; -2.13) | 0.75 (0.34; 1.16) | 0.17 (-0.23; 0.56) | 0.54 (0.34; 0.73) |

**Supplementary Table 4. Estimated coefficients for models of predatory fish dominance (N = 3491).** Based on full fitted models, including main effects of open sea stickleback densities, distance from the open sea, $log_{10}$-transformed wave exposure, an interaction between open sea stickleback densities and distance from the open sea, predation, fishing, temperature, connectivity, interactions between connectivity and predation, between connectivity and fishing, and between temperature and distance from the open sea, as well as a random effect of year. The different representations of connectivity are shown in different columns (3.2 or 3.5 cut-off for wave exposure, distance-weighted sum of all available habitat within a 10 km radius vs network representation). Values in parentheses indicate 95% confidence limits. Values are coloured in green if lower and upper limit both indicate a positive effect, and red if lower and upper limit both indicate a negative effect. Values in bold indicate that effect was included in all candidate models with ΔAIC <4 (indicates substantial support; Burnham & Anderson, 2002). Estimates are based on scaled variables.

| Variable | Connectivity representation | | | |
|---|---|---|---|---|
| | available habitat 3.5 | available habitat 3.2 | network 3.5 | network 3.2 |
| distance to open sea | **1.35 (1.19; 1.51)** | **1.41 (1.25; 1.57)** | **1.4 (1.24; 1.56)** | **1.44 (1.27; 1.60)** |
| open sea stickleback | **-0.74 (-0.88; -0.59)** | **-0.70 (-0.85; -0.55)** | **-0.69 (-0.83; -0.55)** | **-0.69 (-0.83; -0.55)** |
| distance × stickleback | 0.18 (0.03; 0.33) | **0.19 (0.04; 0.34)** | **0.2 (0.05; 0.35)** | **0.20 (0.05; 0.35)** |
| $log_{10}$(wave exposure) | **-0.25 (-0.35; -0.15)** | **-0.28 (-0.39; -0.18)** | **-0.23 (-0.33; -0.13)** | **-0.27 (-0.38; -0.17)** |
| connectivity | **0.16 (0.05; 0.28)** | **0.03 (-0.09; 0.16)** | 0.16 (0.04; 0.27) | **-0.05 (-0.20; 0.10)** |
| predation | **-0.16 (-0.27; -0.04)** | **-0.17 (-0.28; -0.06)** | **-0.16 (-0.27; -0.04)** | **-0.15 (-0.27; -0.04)** |
| fishing | **-0.08 (-0.21; 0.06)** | **-0.14 (-0.29; 0.01)** | **-0.15 (-0.33; 0.03)** | **-0.3 (-0.51; -0.09)** |
| predation × connectivity | **-0.22 (-0.34; -0.11)** | **-0.15 (-0.25; -0.05)** | **-0.23 (-0.34; -0.13)** | **-0.15 (-0.26; -0.05)** |
| fishing × connectivity | 0.20 (0.02; 0.38) | 0.10 (-0.13; 0.33) | 0 (-0.23; 0.22) | -0.21 (-0.52; 0.11) |
| temperature | **0.22 (0.07; 0.38)** | **0.22 (0.07; 0.37)** | **0.20 (0.05; 0.35)** | **0.20 (0.05; 0.35)** |
| temperature × distance | -0.15 (-0.30; 0.00) | -0.17 (-0.32; -0.02) | -0.18 (-0.33; -0.03) | -0.18 (-0.33; -0.03) |

**Supplementary Table 5. Estimated coefficients for model of predatory fish densities (N = 7415).** Based on full fitted models, including main effects of open sea stickleback densities, distance from the open sea, $log_{10}$-transformed wave exposure, an interaction between open sea stickleback densities and distance from the open sea, predation, fishing, temperature, connectivity, interactions between connectivity and predation, between connectivity and fishing, and between temperature and distance from the open sea, as well as a random effect of year. The different representations of connectivity are shown in different columns (3.2 or 3.5 cut-off for wave exposure, distance-weighted sum of all available habitat within a 10 km radius vs network representation). Values in parentheses indicate 95% confidence limits. Values are coloured in green if lower and upper limit both indicate a positive effect, and red if lower and upper limit both indicate a negative effect. Values in bold indicate that effect was included in all candidate models with ΔAIC <4 (indicates substantial support; Burnham & Anderson, 2002). Estimates are based on scaled variables.

| Variable | Connectivity representation | | | |
|---|---|---|---|---|
| | available habitat 3.5 | available habitat 3.2 | network 3.5 | network 3.2 |
| distance to open sea | **0.93 (0.79; 1.06)** | **0.94 (0.82; 1.06)** | **0.93 (0.8; 1.05)** | **0.89 (0.76; 1.01)** |
| open sea stickleback | **0.05 (-0.08; 0.17)** | 0.05 (-0.07; 0.18) | 0.05 (-0.08; 0.17) | 0.03 (-0.1; 0.15) |
| distance × stickleback | **0.22 (0.08; 0.35)** | 0.19 (0.05; 0.33) | 0.22 (0.08; 0.36) | 0.16 (0.02; 0.30) |
| $log_{10}$ (wave exposure) | **-0.23 (-0.34; -0.12)** | **-0.25 (-0.36; -0.13)** | **-0.25 (-0.36; -0.14)** | **-0.26 (-0.37; -0.15)** |
| connectivity | **0.05 (-0.06; 0.16)** | **-0.04 (-0.15; 0.07)** | **0.05 (-0.06; 0.16)** | **-0.01 (-0.14; 0.12)** |
| predation | **-0.74 (-0.85; -0.62)** | **-0.74 (-0.85; -0.62)** | **-0.67 (-0.78; -0.55)** | **-0.63 (-0.74; -0.52)** |
| fishing | -0.07 (-0.17; 0.03) | -0.09 (-0.21; 0.03) | -0.06 (-0.22; 0.10) | 0.05 (-0.14; 0.24) |
| predation × connectivity | **-0.47 (-0.60; -0.35)** | **-0.47 (-0.57; -0.36)** | **-0.36 (-0.47; -0.25)** | **-0.41 (-0.52; -0.31)** |
| fishing × connectivity | 0.02 (-0.11; 0.14) | -0.02 (-0.20; 0.16) | 0.01 (-0.18; 0.20) | 0.20 (-0.07; 0.46) |
| temperature | **0.29 (0.11; 0.47)** | **0.24 (0.05; 0.42)** | **0.25 (0.06; 0.43)** | **0.22 (0.03; 0.40)** |
| temperature × distance | **-0.25 (-0.39; -0.12)** | **-0.3 (-0.44; -0.17)** | **-0.3 (-0.44; -0.16)** | **-0.33 (-0.47; -0.19)** |

**Supplementary Table 6. Estimated coefficients for model of stickleback densities (N =7167)**. Based on full fitted models, including main effects of open sea stickleback densities, distance from the open sea, $log_{10}$-transformed wave exposure, an interaction between open sea stickleback densities and distance from the open sea, predation, fishing, temperature, connectivity, interactions between connectivity and predation, between connectivity and fishing, and between temperature and distance from the open sea, as well as a random effect of year. The different representations of connectivity are shown in different columns (3.2 or 3.5 cut-off for wave exposure, distance-weighted sum of all available habitat within a 10 km radius vs network representation). Values in parentheses indicate 95% confidence limits. Values are coloured in green if lower and upper limit both indicate a positive effect, and red if lower and upper limit both indicate a negative effect. Values in bold indicate that effect was included in all candidate models with ΔAIC <4 (indicates substantial support; Burnham & Anderson, 2002). Estimates are based on scaled variables.

| Variable | Connectivity representation | | | |
|---|---|---|---|---|
| | available habitat 3.5 | available habitat 3.2 | network 3.5 | network 3.2 |
| distance to open sea | **-1.18 (-1.36; -1.00)** | **-1.2 (-1.37; -1.02)** | **-1.16 (-1.33; -0.98)** | **-1.15 (-1.33; -0.97)** |
| open sea stickleback | **1.18 (0.97; 1.38)** | **1.18 (0.98; 1.39)** | **1.19 (0.99; 1.40)** | **1.20 (1.00; 1.40)** |
| distance × stickleback | **0.50 (0.30; 0.71)** | **0.47 (0.27; 0.67)** | **0.54 (0.34; 0.75)** | **0.51 (0.31; 0.72)** |
| $log_{10}$(wave exposure) | **0.39 (0.24; 0.54)** | **0.40 (0.25; 0.55)** | **0.41 (0.26; 0.56)** | **0.42 (0.27; 0.58)** |
| connectivity | -0.10 (-0.24; 0.05) | -0.08 (-0.24; 0.08) | 0.03 (-0.13; 0.19) | **-0.01 (-0.20; 0.19)** |
| predation | **-0.32 (-0.45; -0.19)** | **-0.32 (-0.45; -0.19)** | **-0.34 (-0.48; -0.21)** | **-0.32 (-0.46; -0.19)** |
| fishing | -0.14 (-0.29; 0.01) | -0.16 (-0.33; 0.01) | -0.10 (-0.36; 0.15) | -0.30 (-0.56; -0.03) |
| predation × connectivity | 0.00 (-0.15; 0.15) | -0.11 (-0.24; 0.01) | 0.03 (-0.11; 0.18) | -0.03 (-0.16; 0.11) |
| fishing × connectivity | -0.28 (-0.44; -0.12) | -0.31 (-0.54; -0.07) | -0.14 (-0.45; 0.17) | -0.45 (-0.83; -0.07) |
| temperature | **0.16 (-0.09; 0.40)** | 0.13 (-0.11; 0.37) | **0.16 (-0.08; 0.40)** | **0.09 (-0.15; 0.34)** |
| temperature × distance | **0.31 (0.10; 0.52)** | 0.28 (0.07; 0.48) | **0.35 (0.16; 0.55)** | **0.36 (0.16; 0.57)** |

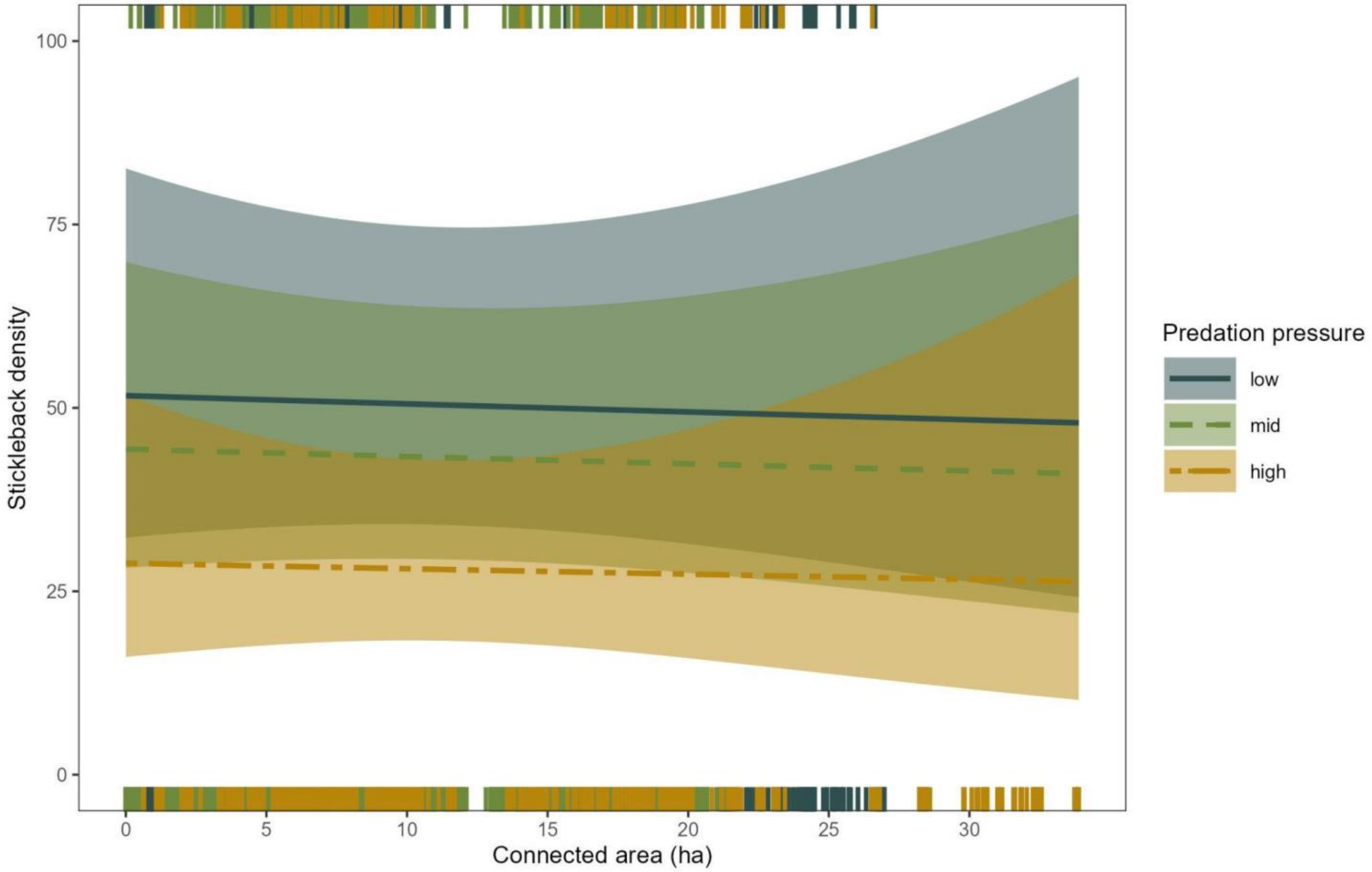


**Supplementary Fig. 3. Predicted stickleback density, as a function of connectivity for different levels of predation pressure (low = 10th percentile, mid = median, high = 90th percentile).** Lines show predictions from a generalised linear mixed model also including a random effect of year and main effects of open sea stickleback densities, distance from the open sea, $\log_{10}$-transformed wave exposure, an interaction between open sea stickleback densities and distance from the open sea, fishing, temperature, interactions between connectivity and fishing, and between temperature and distance from the open sea, with 95% confidence intervals. The connectivity measure used is distance-weighted sum of all available habitat within a 10 km radius with a 3.5 cut-off for wave exposure. Stickleback densities refer to the number of individuals in a sampling area of roughly 80 $m^2$.

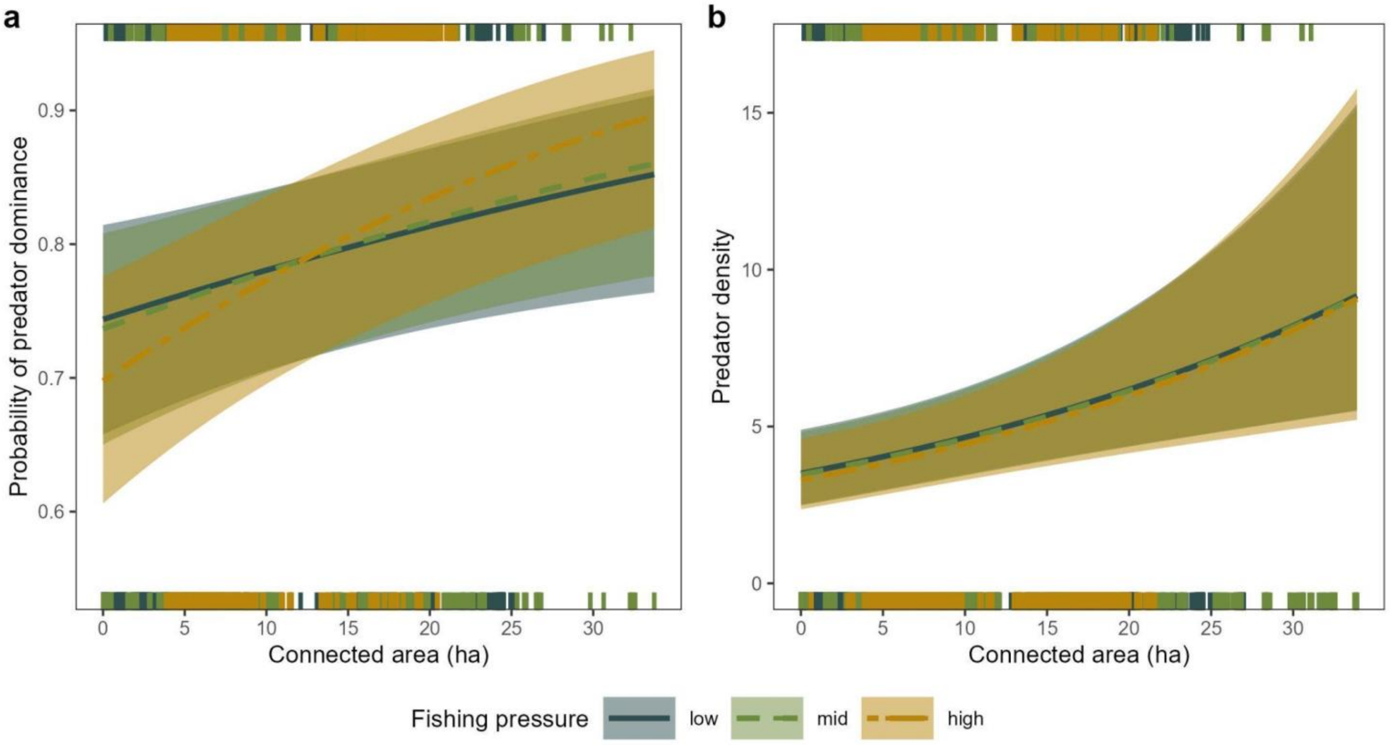


**Supplementary Fig. 4. Predicted (a) relative dominance of predatory fish (0 = stickleback dominance, 1 = predatory fish dominance) and (b) predatory fish density, as a function of connectivity for different levels of fishing pressure (low = 10th percentile, mid = median, high = 90th percentile).** Lines show predictions from a generalised linear mixed model also including a random effect of year and main effects of open sea stickleback densities, distance from the open sea, $\log_{10}$-transformed wave exposure, an interaction between open sea stickleback densities and distance from the open sea, predation, temperature, an interaction between connectivity and predation, and an interaction between temperature and distance from the open sea, with 95% confidence intervals. The connectivity measure used is distance-weighted sum of all available habitat within a 10 km radius with a 3.5 cut-off for wave exposure. Predator densities refer to the number of individuals in a sampling area of roughly 80 $m^2$.

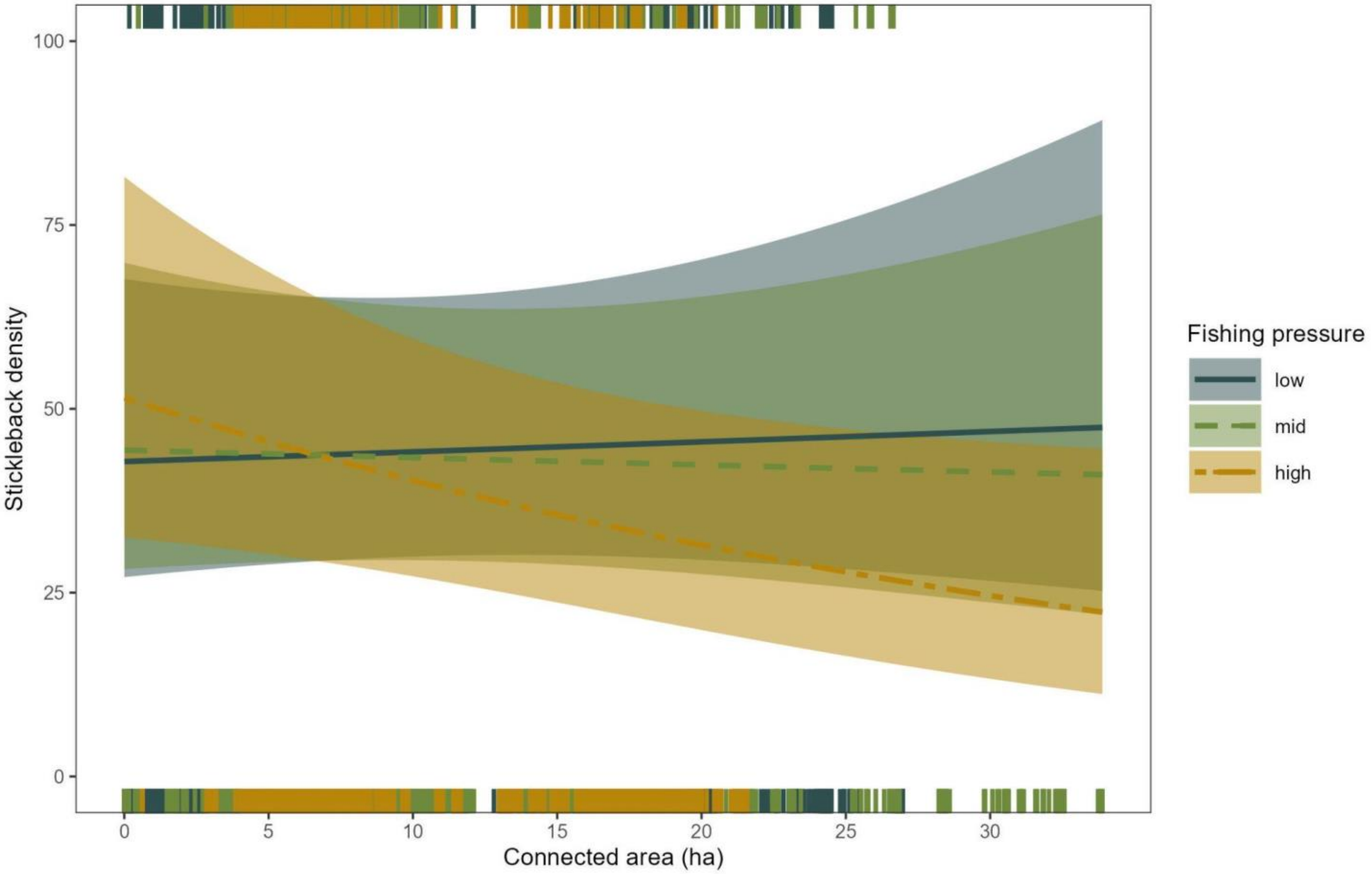


**Supplementary Fig. 5. Predicted stickleback density, as a function of connectivity for different levels of fishing pressure (low = 10th percentile, mid = median, high = 90th percentile).** Lines show predictions from a generalised linear mixed model also including a random effect of year and main effects of open sea stickleback densities, distance from the open sea, $\log_{10}$-transformed wave exposure, an interaction between open sea stickleback densities and distance from the open sea, predation, temperature, interactions between connectivity and predation, and between temperature and distance from the open sea, with 95% confidence intervals. The connectivity measure used is distance-weighted sum of all available habitat within a 10 km radius with a 3.5 cut-off for wave exposure. Stickleback densities refer to the number of individuals in a sampling area of roughly 80 $m^2$.

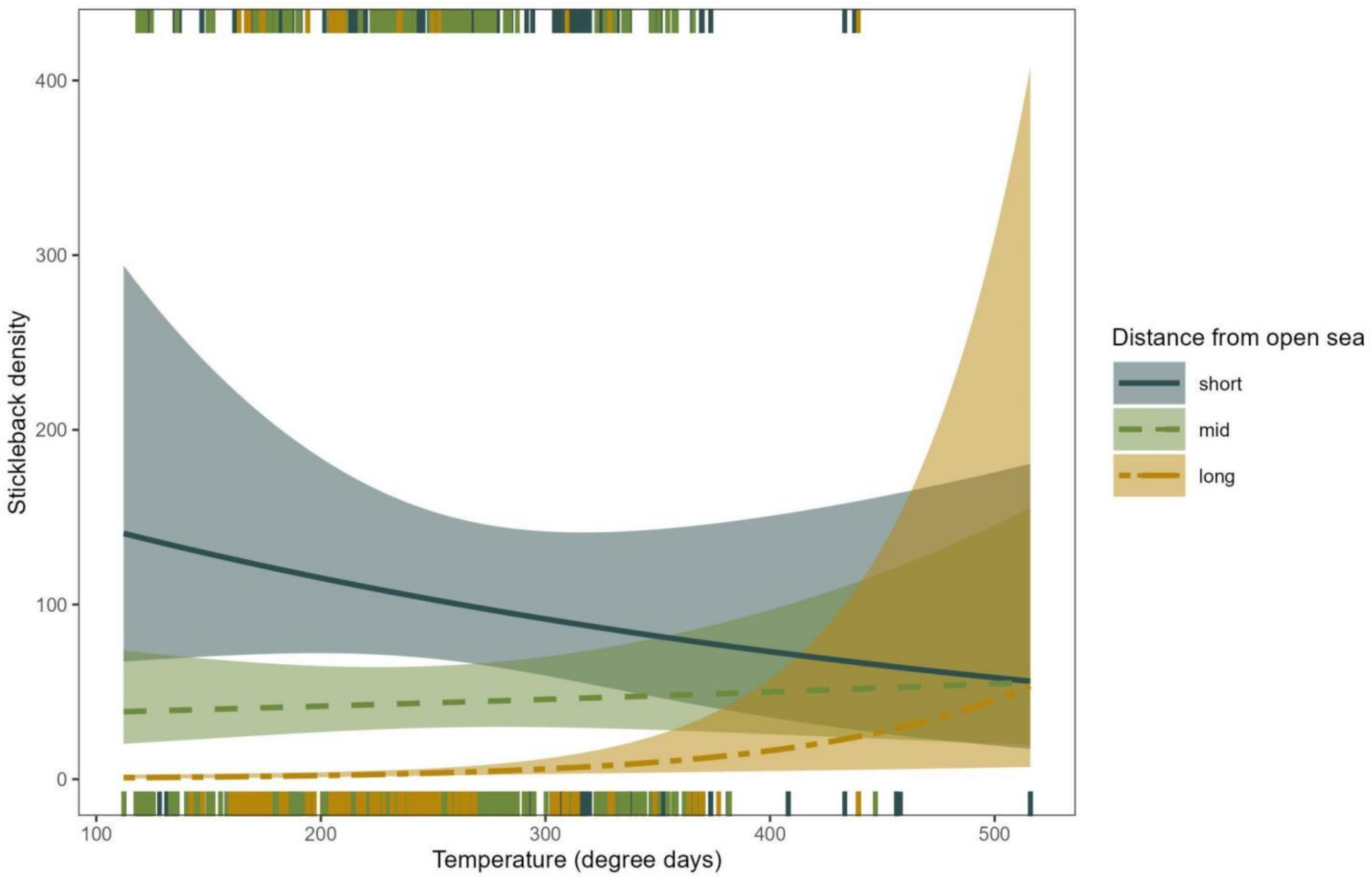


**Supplementary Fig. 6. Predicted stickleback density, as a function of temperature for different distances from the open sea (short = 10th percentile, mid = median, long = 90th percentile).** Lines show predictions from a generalised linear mixed model also including a random effect of year and main effects of open sea stickleback densities, $\log_{10}$-transformed wave exposure, an interaction between open sea stickleback densities and distance from the open sea, predation, fishing, and interactions between connectivity and predation and between connectivity and fishing with 95% confidence intervals. The connectivity measure used is distance-weighted sum of all available habitat within a 10 km radius with a 3.5 cut-off for wave exposure. Stickleback densities refer to the number of individuals in a sampling area of roughly 80 $m^2$.

**Supplementary Table 7. Comparison of models and connectivity representations for predatory fish dominance, predatory fish densities and stickleback densities.** The baseline model includes main effects of distance from the open sea, offshore stickleback densities, $log_{10}$-transformed wave exposure and an interaction between distance from the open sea and offshore stickleback. The “baseline + resilience” models include main effects of open sea stickleback densities, distance from the open sea, $log_{10}$-transformed wave exposure, an interaction between open sea stickleback densities and distance from the open sea, predation, fishing, temperature, connectivity, interactions between connectivity and predation, between connectivity and fishing, and between temperature and distance from the open sea. For each of these models, the type of connectivity metric used is indicated (3.2 or 3.5 cut-off for wave exposure, distance-weighted sum of all available habitat within a 10 km radius vs network representation). Finally, the “resilience only” model includes main effects of predation, fishing, temperature, connectivity, interactions between connectivity and predation, between connectivity and fishing, and between temperature and distance from the open sea. Here, the distance-weighted sum of all available habitat within a 10 km radius with a 3.5 cut-off for wave exposure is used as this was the connectivity representation that showed the best cross-model performance. All models include a random effect of year. Shows $R^2$ calculated using the function *r.squaredGLMM* from the package MuMIn (Bartoń, 2020) using the delta method, as well as ΔAIC relative to the baseline model, also calculated using the MuMIn-package.

| Model | Predatory fish dominance | | Predatory fish density | | Stickleback density | |
|---|---|---|---|---|---|---|
| | $R^2$ | ΔAIC | $R^2$ | ΔAIC | $R^2$ | ΔAIC |
| Baseline | 0.34 | 0.00 | 0.23 | 0.00 | 0.36 | 0.00 |
| Baseline + resilience available habitat 3.5 | 0.38 | -52.76 | 0.38 | -281.06 | 0.38 | -23.57 |
| Baseline + resilience available habitat 3.2 | 0.37 | -35.30 | 0.38 | -299.08 | 0.38 | -23.19 |
| Baseline + resilience network 3.5 | 0.37 | -61.80 | 0.36 | -260.70 | 0.38 | -15.76 |
| Baseline + resilience network 3.2 | 0.37 | -39.78 | 0.36 | -280.97 | 0.38 | -24.64 |
| Resilience only network 3.5 | 0.31 | 89.34 | 0.36 | -260.16 | 0.25 | 108.42 |

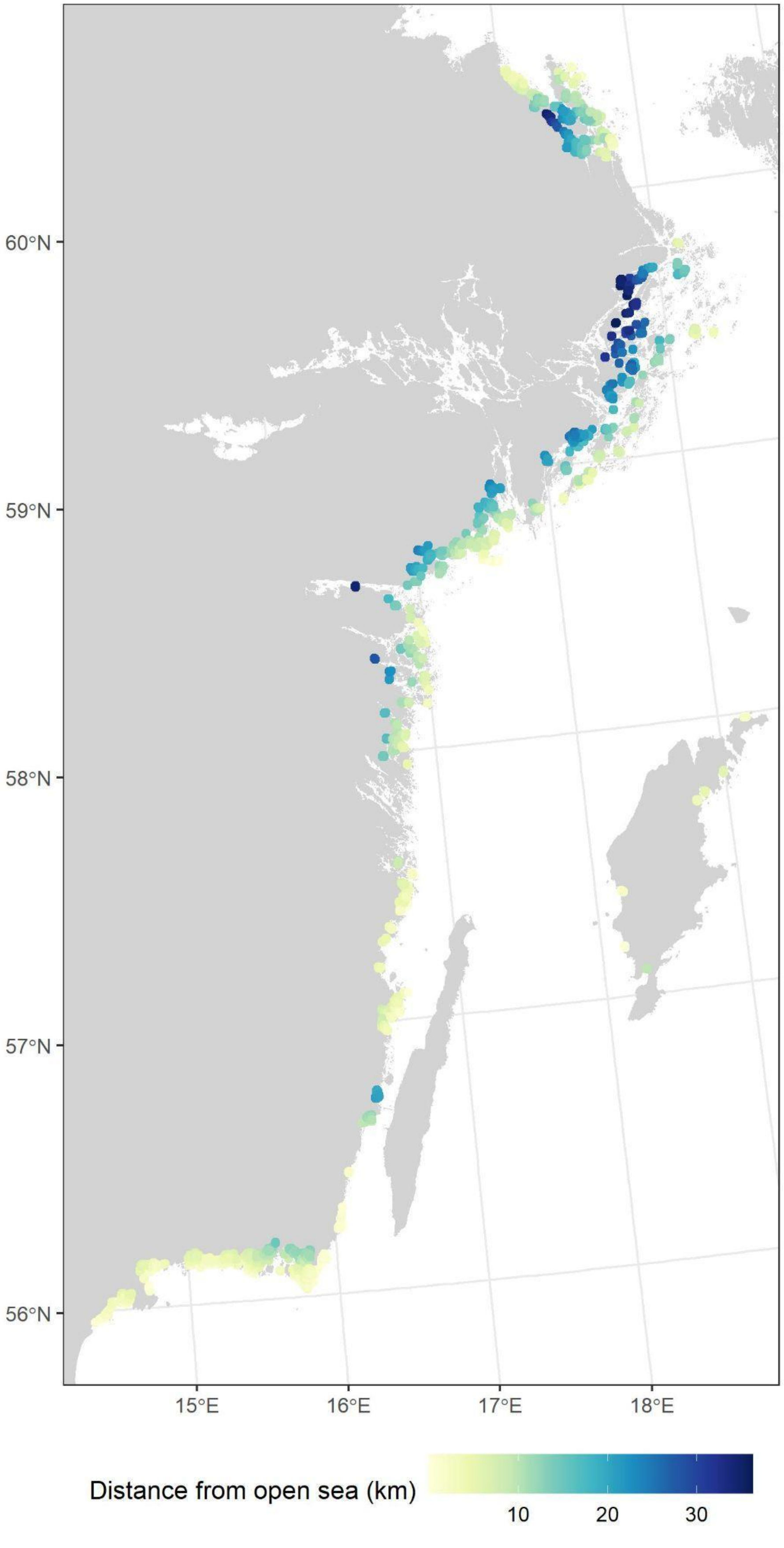


**Supplementary Fig. 7. Map of extracted distances from the open sea for each detonation point.**

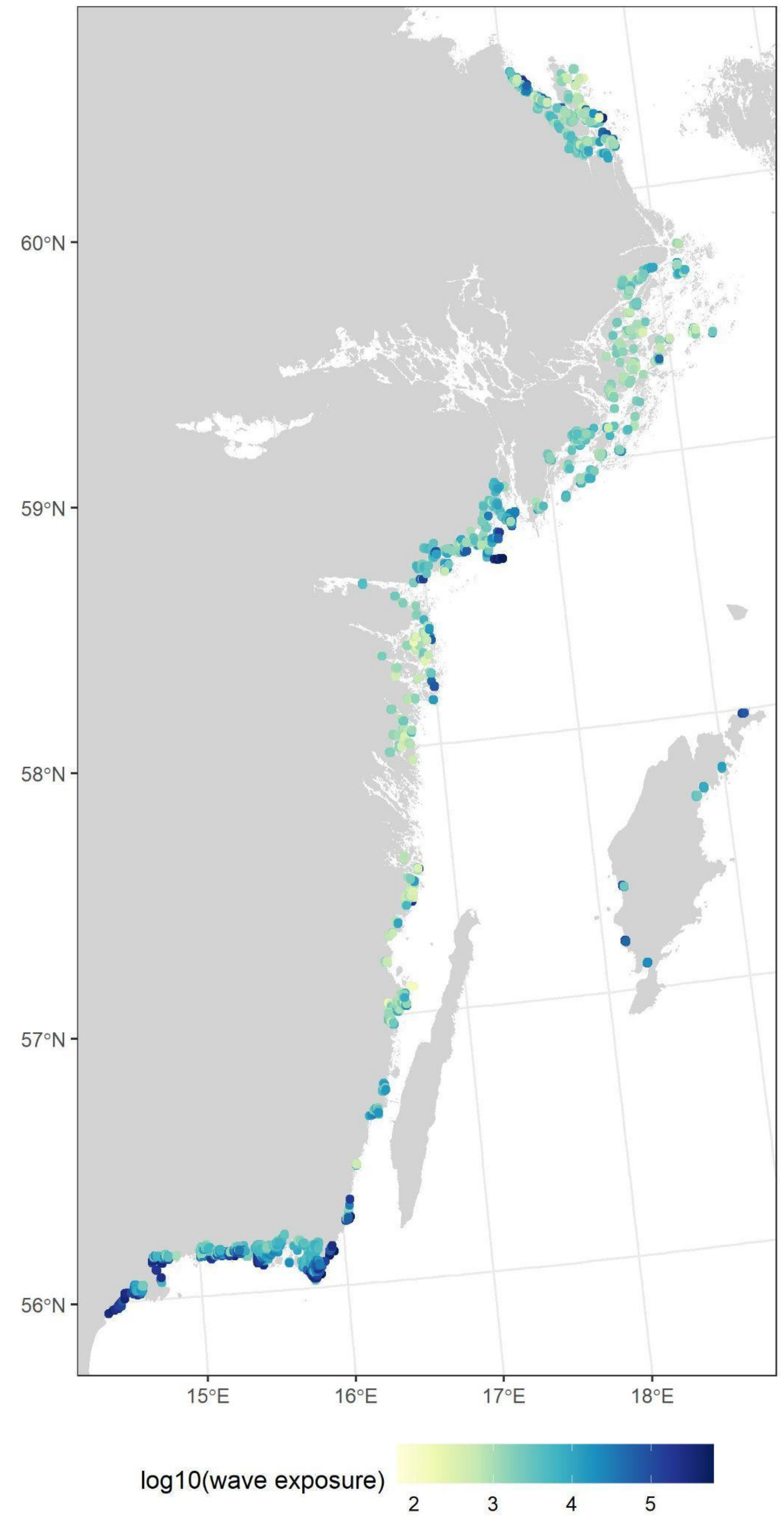


**Supplementary Fig. 8. Map of extracted values of $\log_{10}$-transformed wave exposure for each detonation point.**

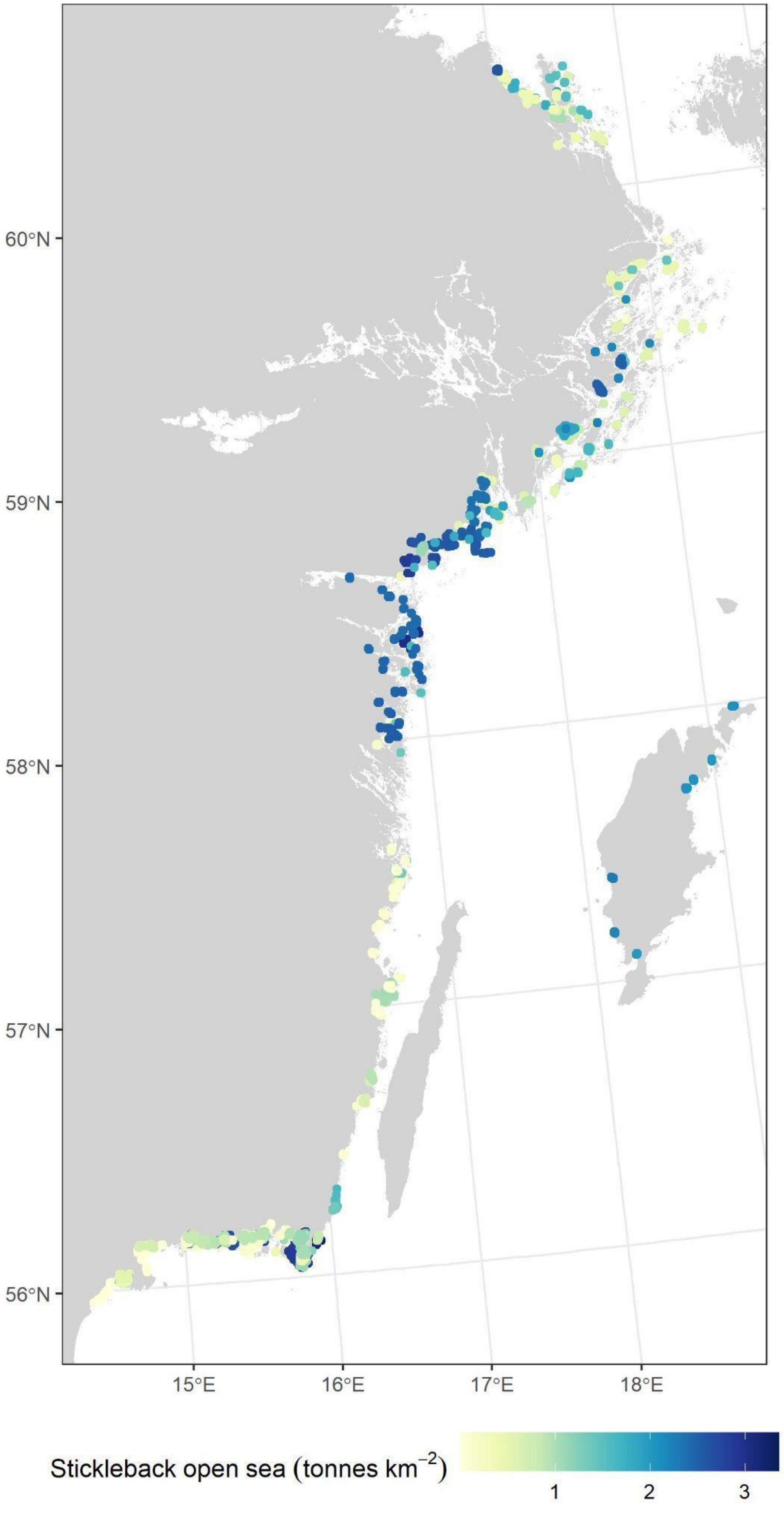


**Supplementary Fig. 9**. **Map of extracted values of open sea stickleback densities for each detonation point.** Note that this varies between years, which is why close-by locations can have very different values.

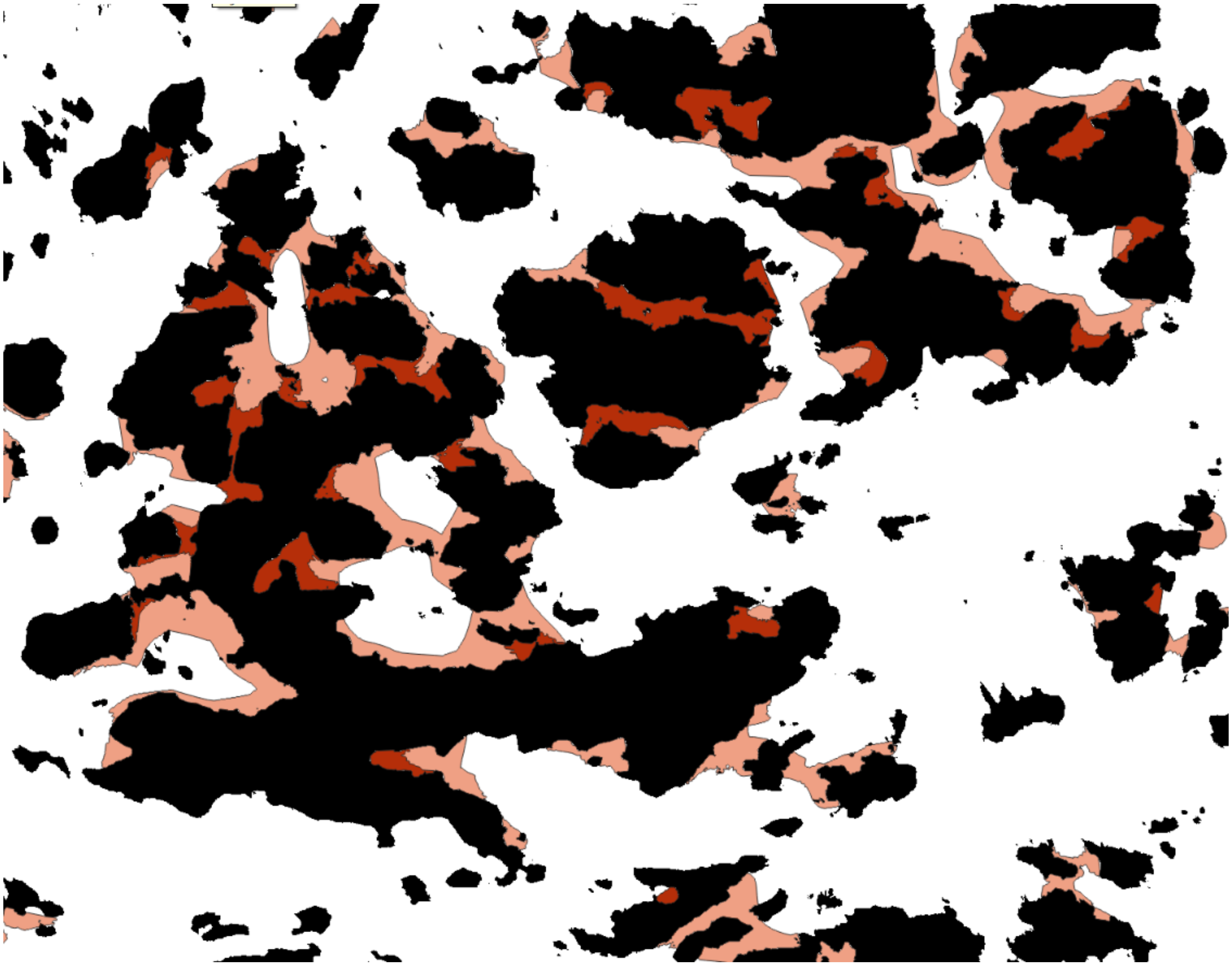

**Supplementary Fig. 10. Illustration of difference in habitat extent between the 3.2 and 3.5 cut-off for log$_{10}$-transformed wave exposure**. Black = land, light red = 3.5 cut-off, and darker red = 3.2 cut-off. Note that all habitat included in the 3.2 cut-off habitat is also included in the 3.5 cut-off habitat. The reason for using two cut-off levels for wave exposure was that this cut-off has a large impact on the extent of predicted habitat (as seen in this image), while also being uncertain and potentially varying between years.

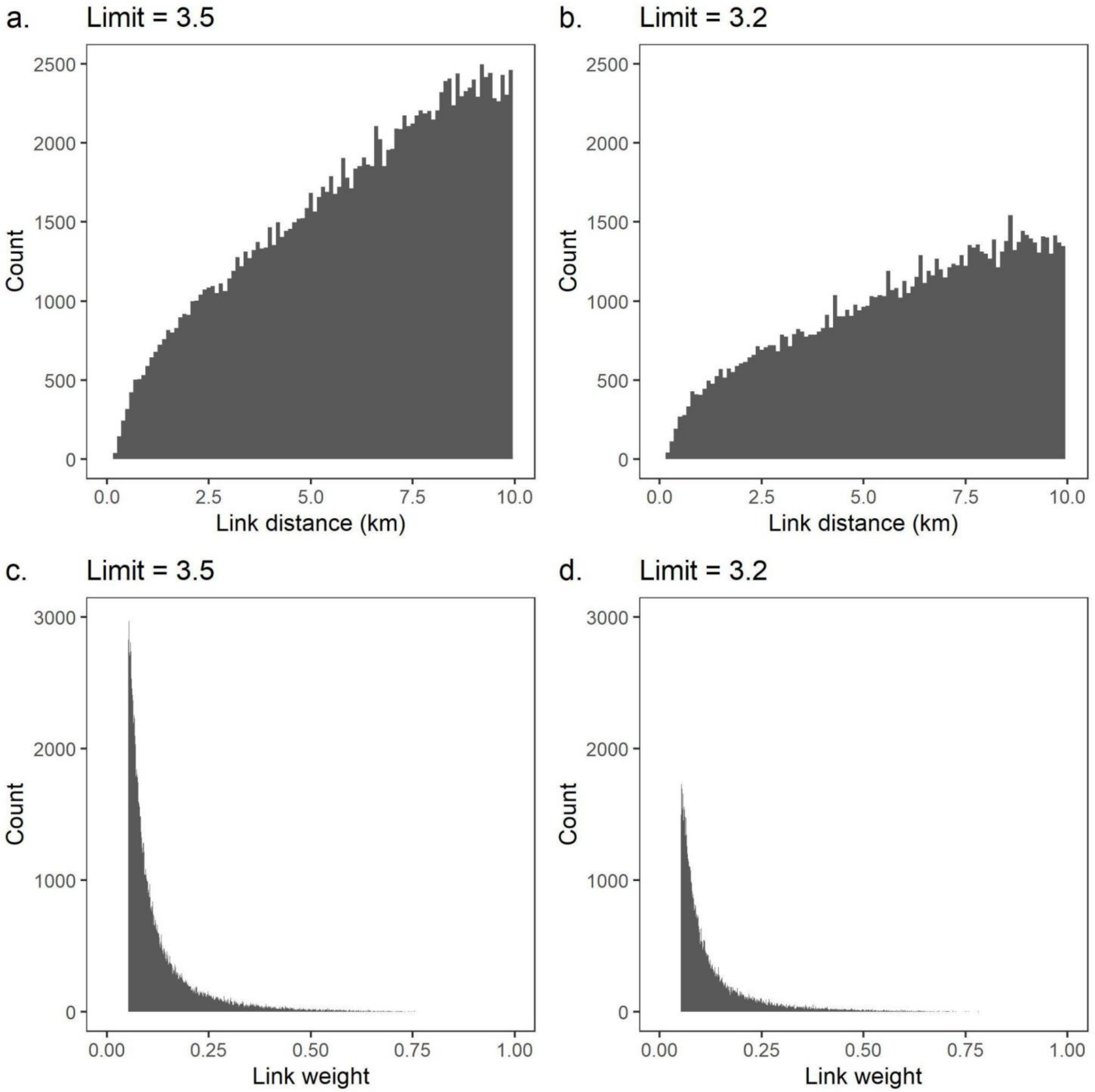


**Supplementary Fig. 11. Distribution of link distances (a,b) and corresponding link weights (c,d) based on a 3.5 cut-off (a,c), and a 3.2 cut-off (b,d) for $log_{10}$-transformed wave exposure.** Note that most patch pairs are connected by longer distances (and thus weakly connected). Note also that we used 10 km as cut-off, which is why link weight does not quite reach zero.

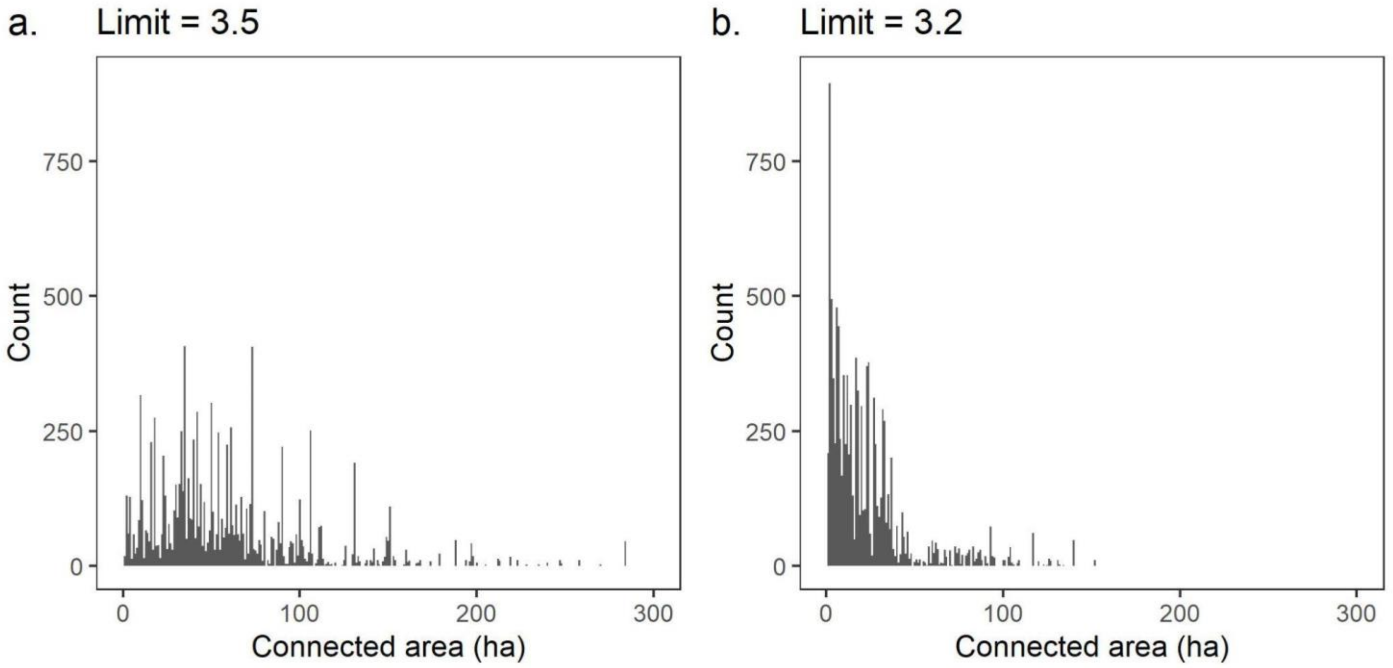


**Supplementary Fig. 12. Histogram of calculated connected areas for all detonation points based on a network representation of connectivity and (a) a 3.5 cut-off, and (b) a 3.2 cut-off for $log_{10}$-transformed wave exposure.**

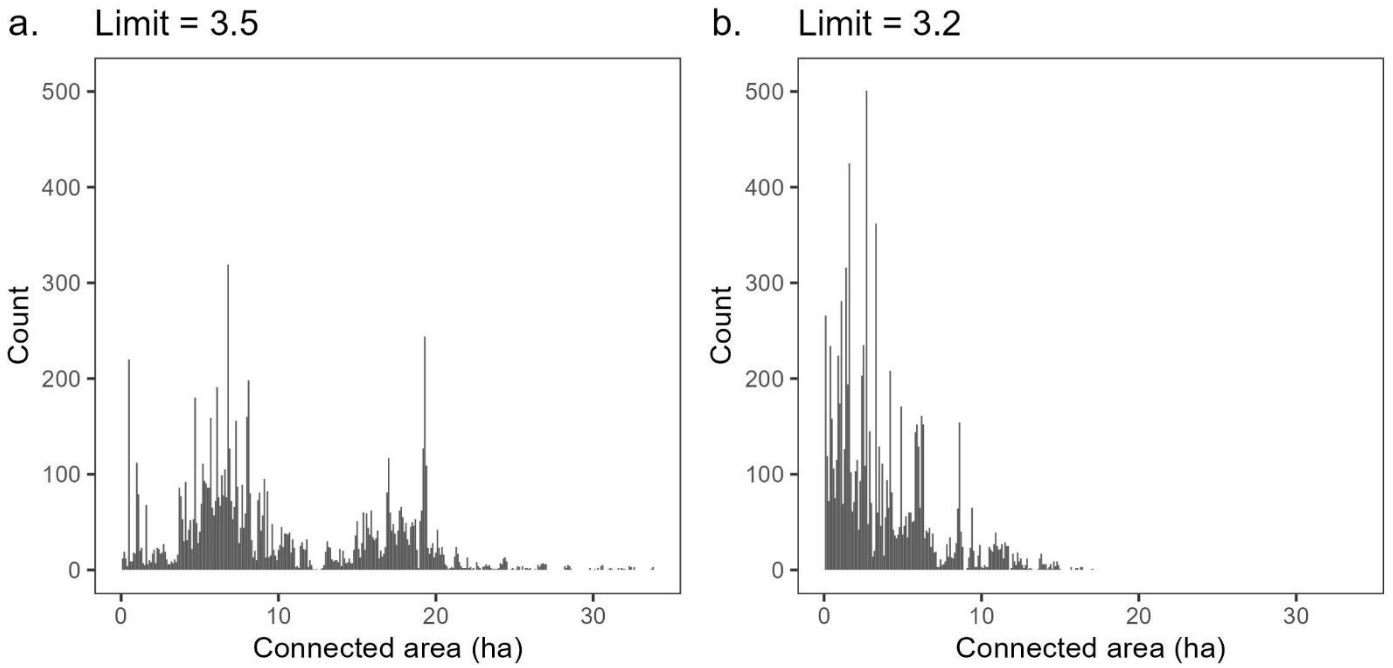


**Supplementary Fig. 13. Histogram of calculated connected area for all detonation points based on a distance-weighted sum of all available habitat within a 10 km radius representation of connectivity and (a) a 3.5 cut-off, and (b) a 3.2 cut-off for $log_{10}$-transformed wave exposure.**

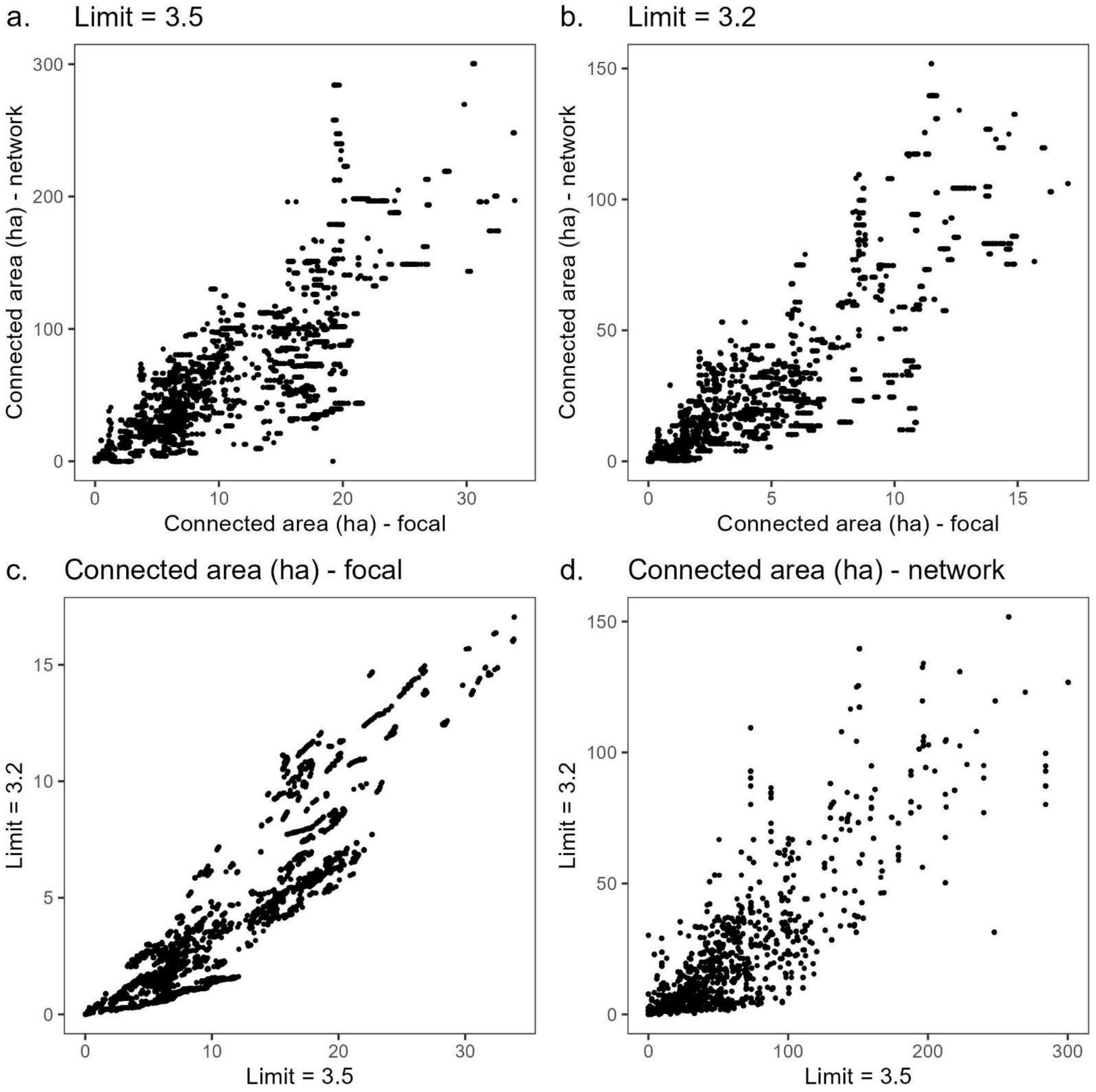


**Supplementary Fig. 14. Comparison of calculated connectivity values for the different approaches.** (a) network representation (network) vs distance-weighted sum of all available habitat within a 10 km radius (focal) for a 3.5 cut-off for $log_{10}$-transformed wave exposure, (b) network representation (network) vs distance-weighted sum of all available habitat within a 10 km radius (focal) for a 3.2 cut-off for $log_{10}$-transformed wave exposure, (c) 3.5 vs 3.2 cut-off for $log_{10}$-transformed wave exposure for distance-weighted sum of all available habitat within a 10 km radius (focal) and (d) 3.5 vs 3.2 cut-off for $log_{10}$-transformed wave exposure for the network representation of connectivity (network).

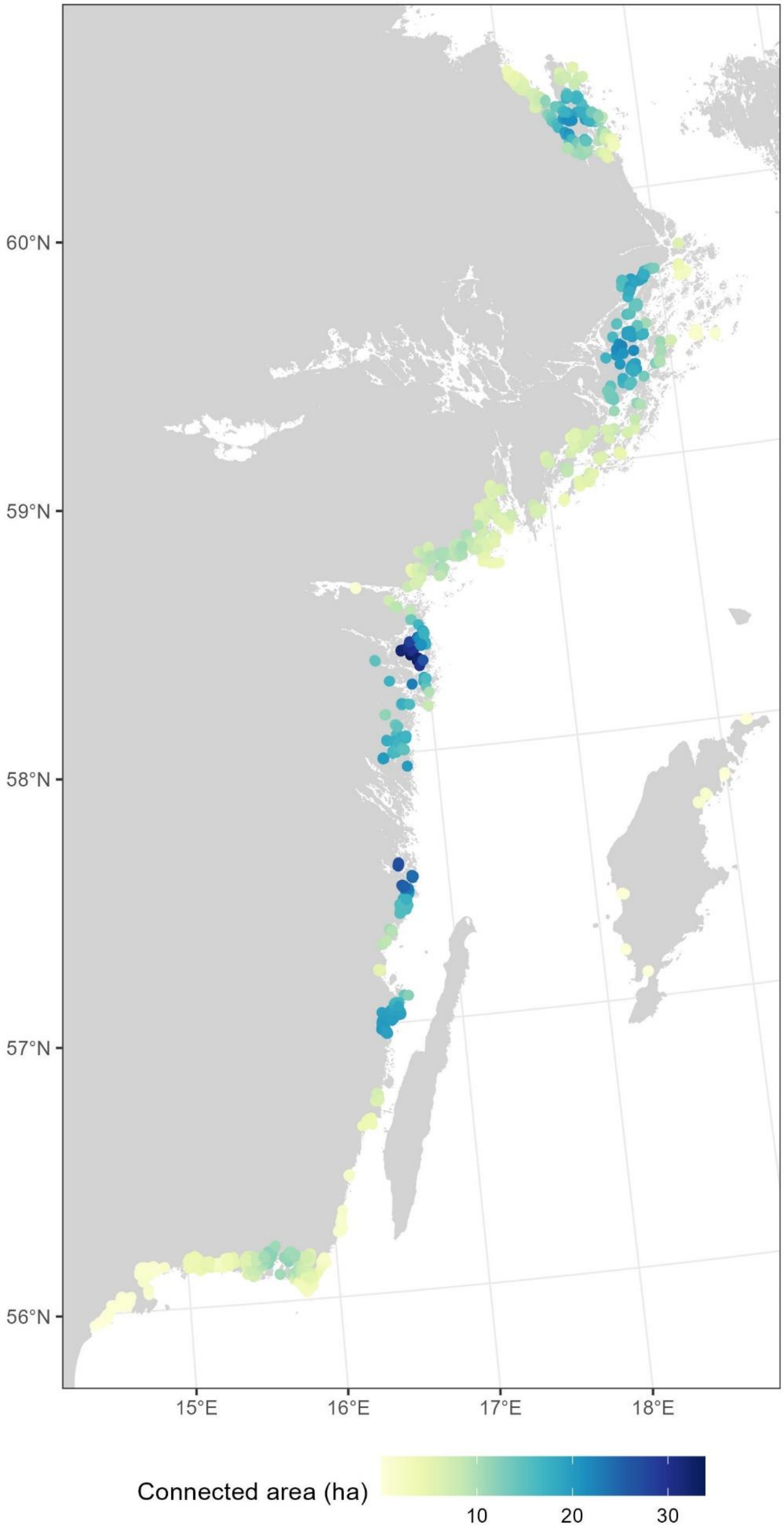


**Supplementary Fig. 15. Map of extracted connectivity for each detonation point, calculated as weighted sum of all available habitat within a 10 km radius and using a 3.5 cut-off for $log_{10}$-transformed wave exposure.**

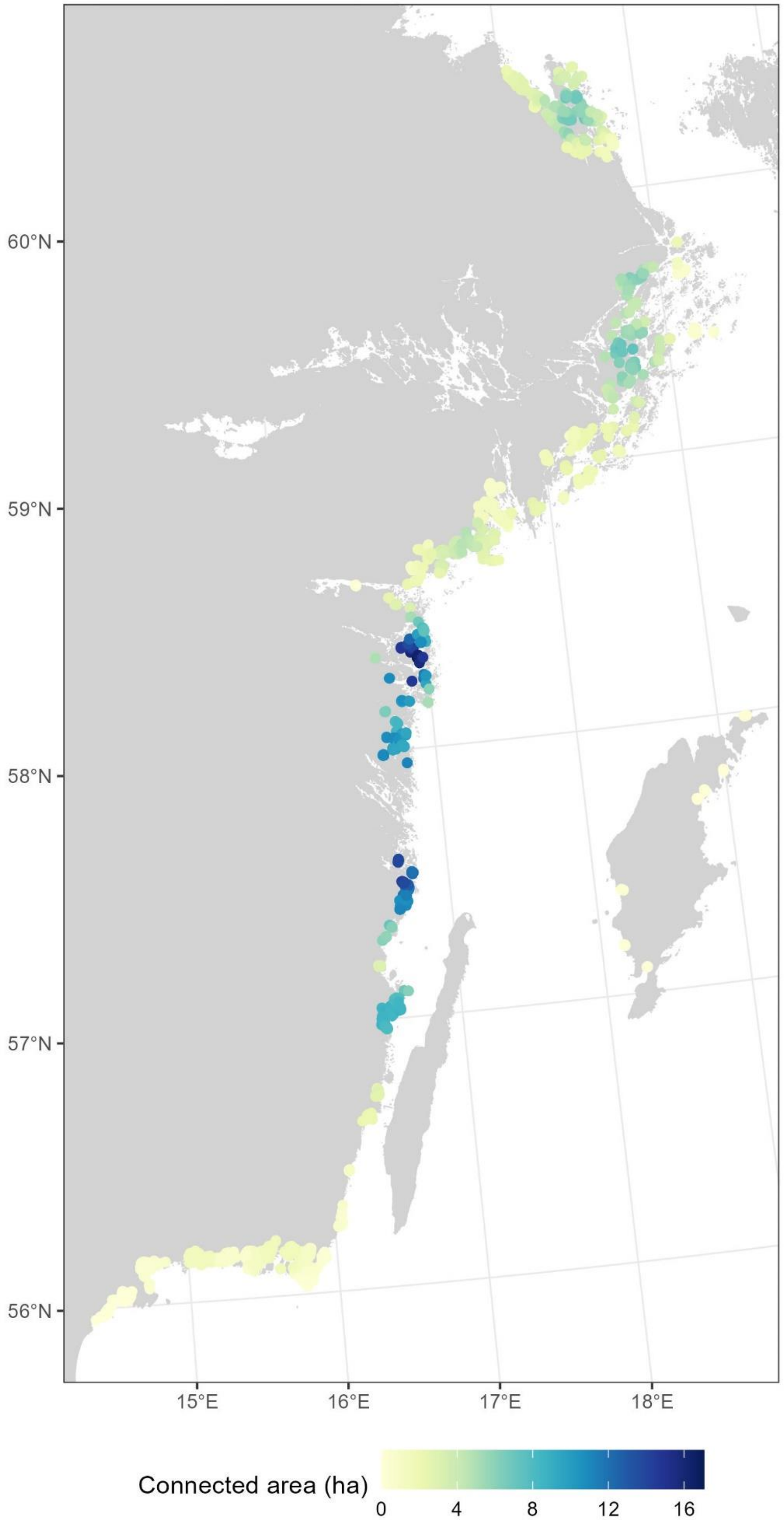


**Supplementary Fig. 16. Map of extracted connectivity for each detonation point, calculated as weighted sum of all available habitat within a 10 km radius and using a 3.2 cut-off for $\log_{10}$-transformed wave exposure.**

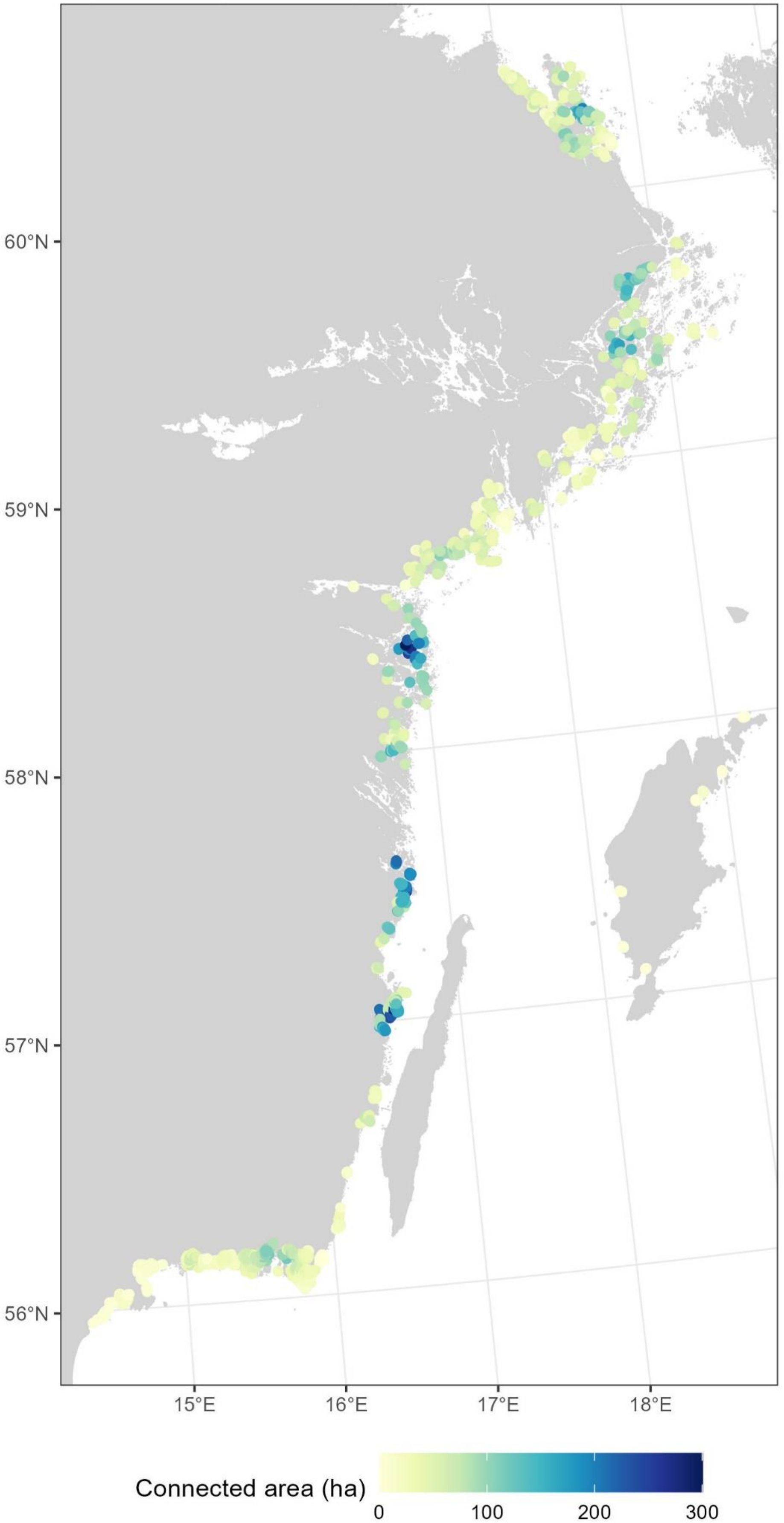


**Supplementary Fig. 17. Map of extracted connectivity for each detonation point, calculated based on the created network and using a 3.5 cut-off for $log_{10}$-transformed wave exposure.**

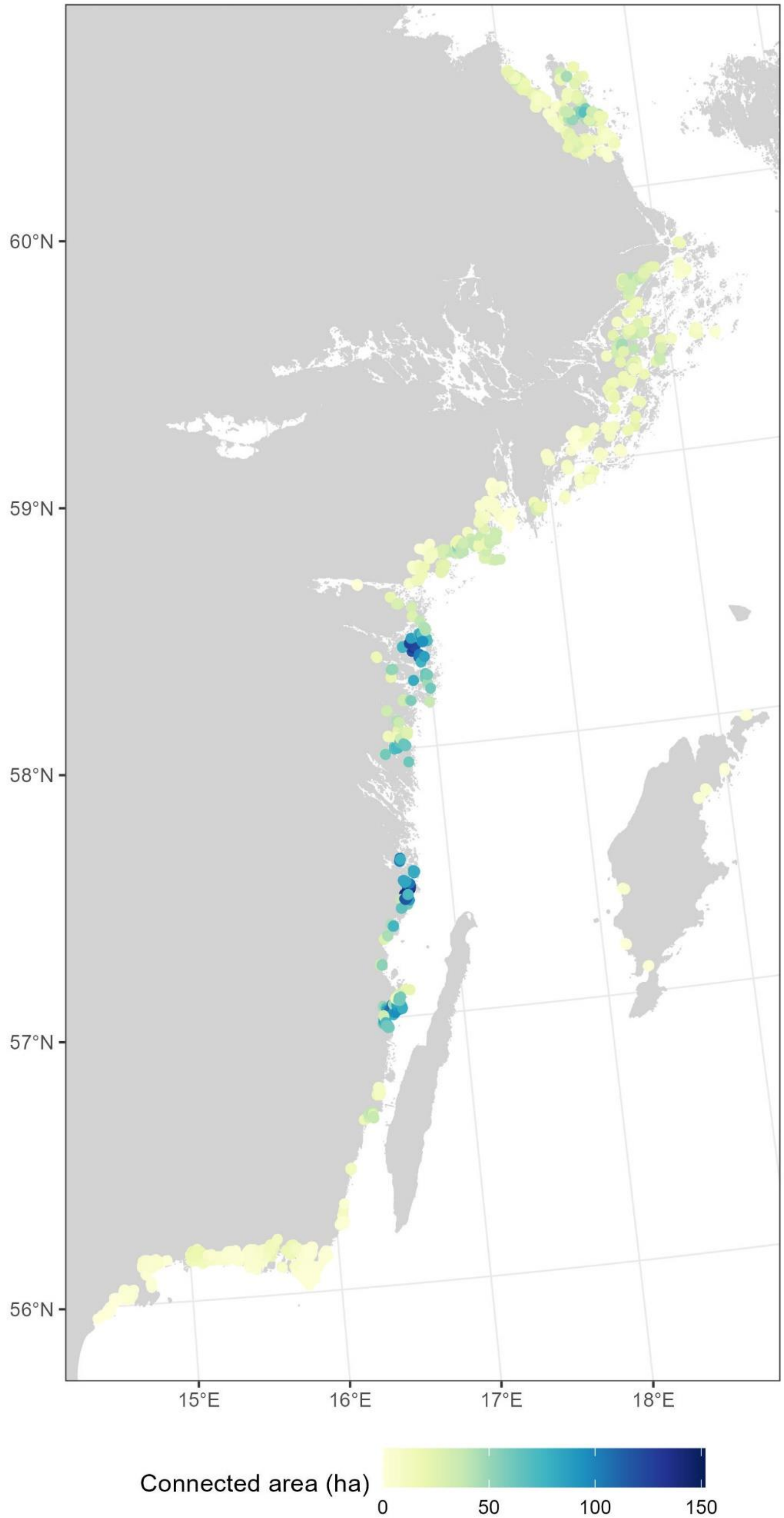


**Supplementary Fig. 18. Map of extracted connectivity for each detonation point, calculated based on the created network and using a 3.2 cut-off for $log_{10}$-transformed wave exposure.**

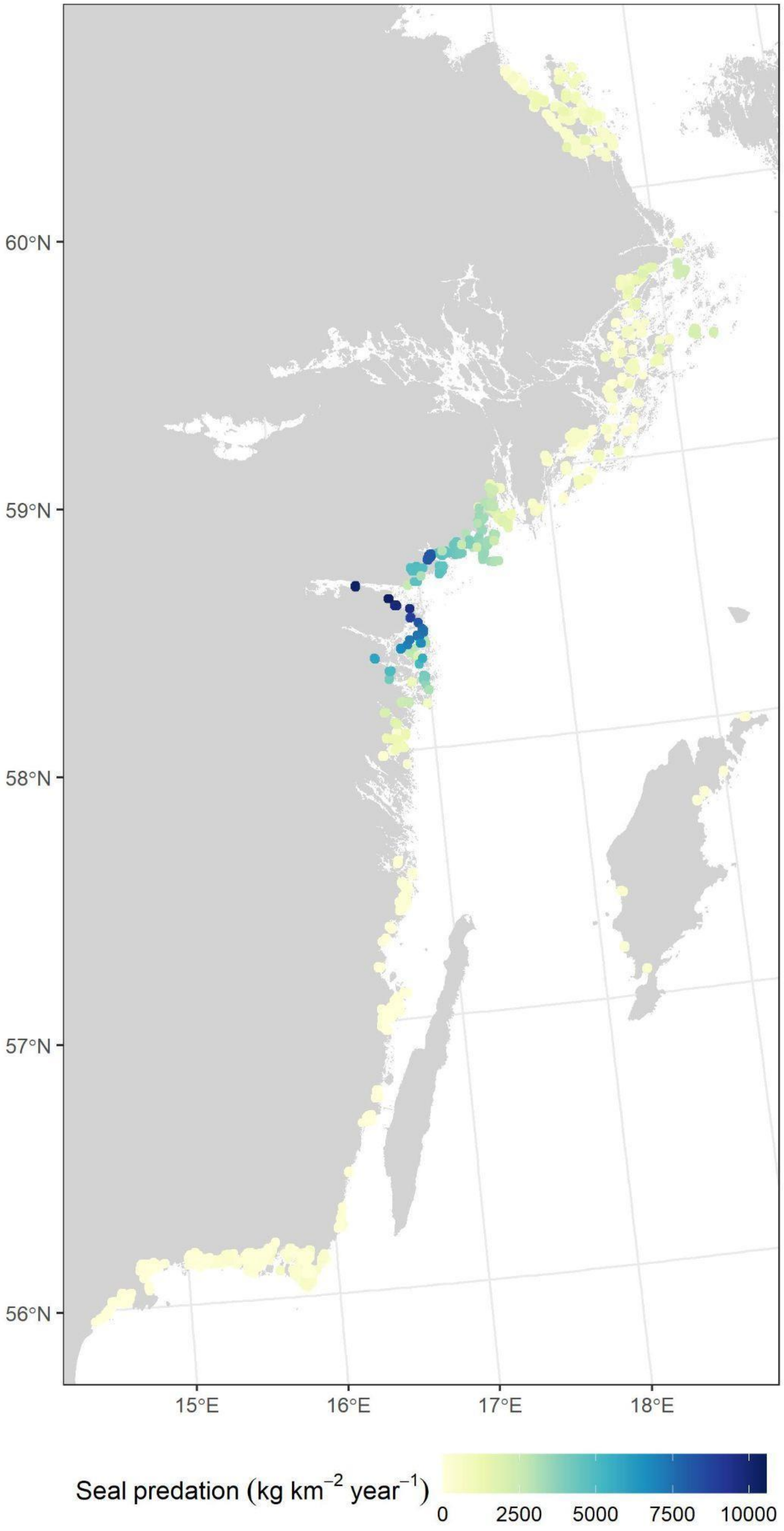


**Supplementary Fig. 19**. **Map of extracted values of seal predation for each detonation point.** Note that this varies between years, which is why close-by locations can have quite different values.

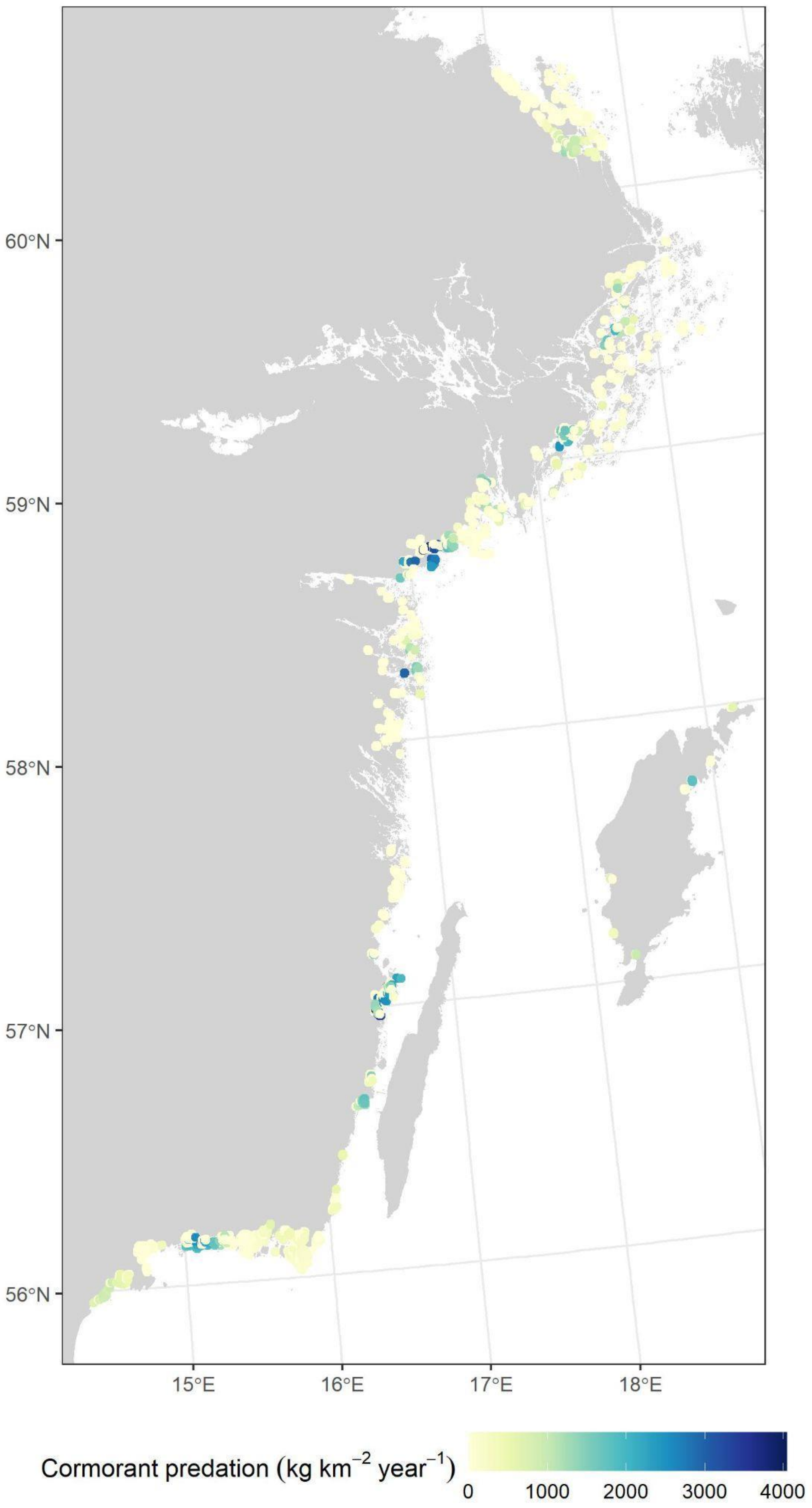


**Supplementary Fig. 20**. **Map of extracted values of cormorant predation for each detonation point.** Note that this varies between years, which is why close-by locations can have very different values.

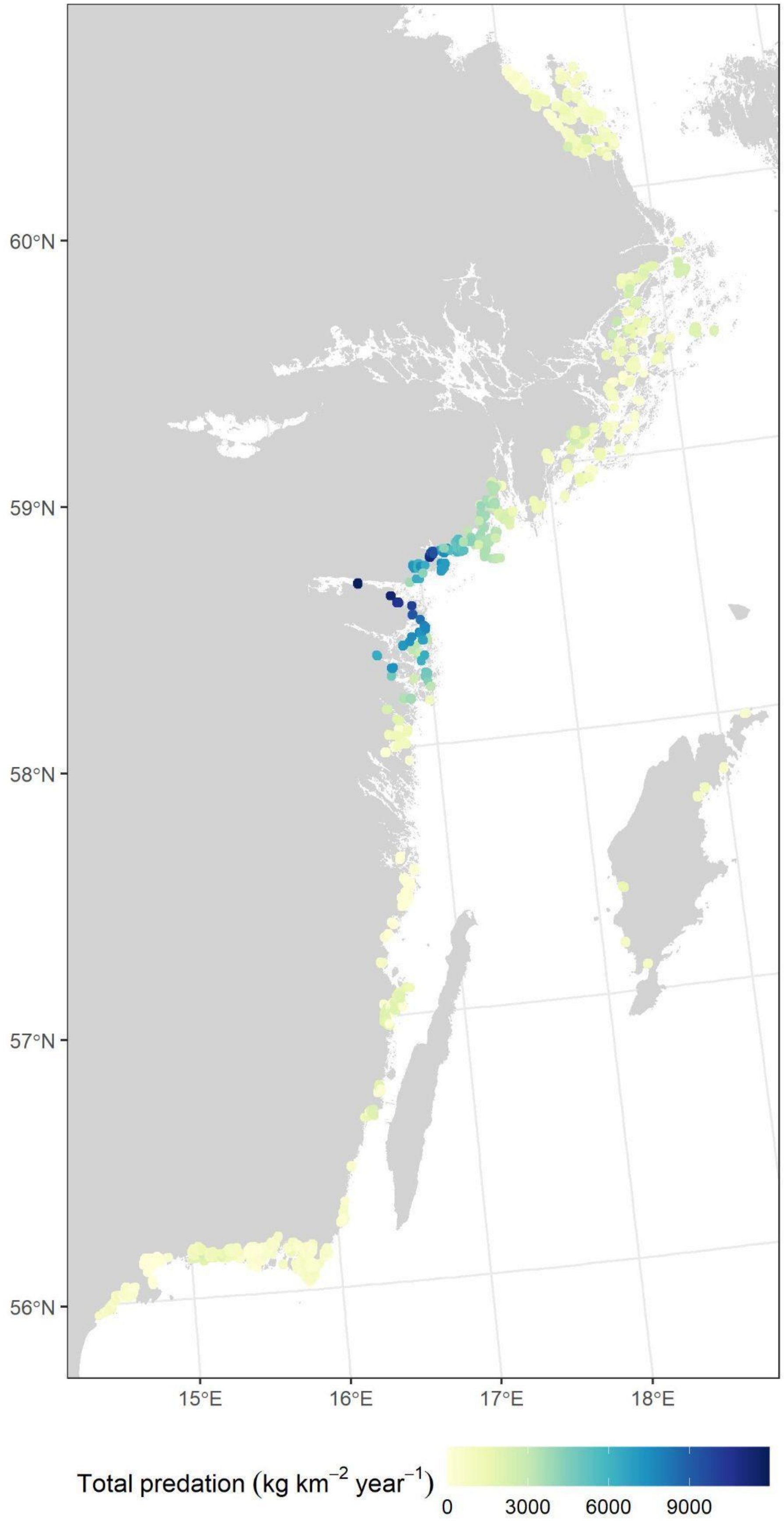


**Supplementary Fig. 21**. **Map of extracted values of seal and cormorant predation added together for each detonation point.** Note that this varies between years, which is why close-by locations can have quite different values.

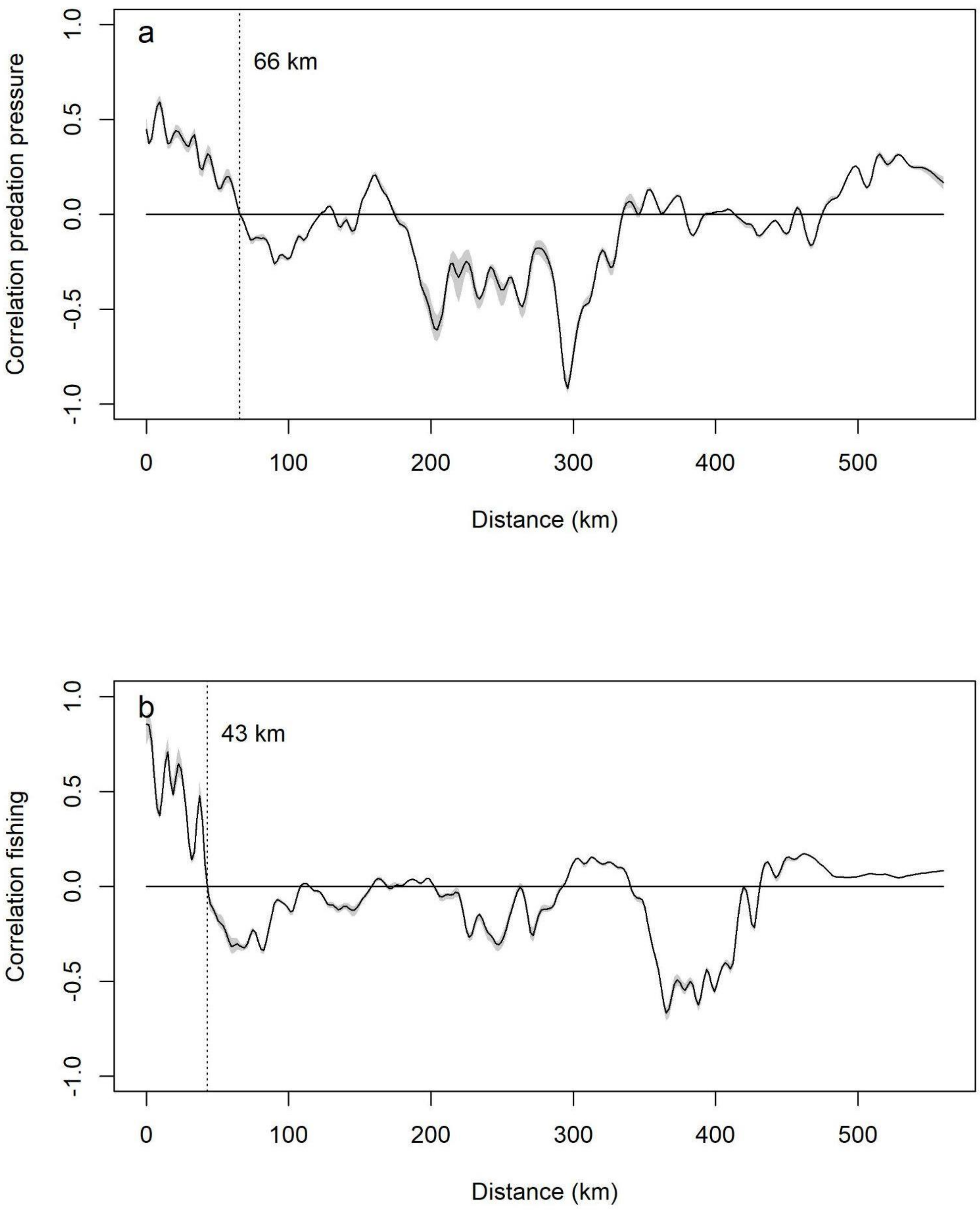


**Supplementary Fig. 22. Spline correlogram of extracted values for (a) predation pressure and (b) fishing pressure.** Dotted lines with indicated distances show where spline crosses 0 for the first time. Produced using the *spline.correlogram*-function in the R-package *ncf* (Bjørnstad, 2022).

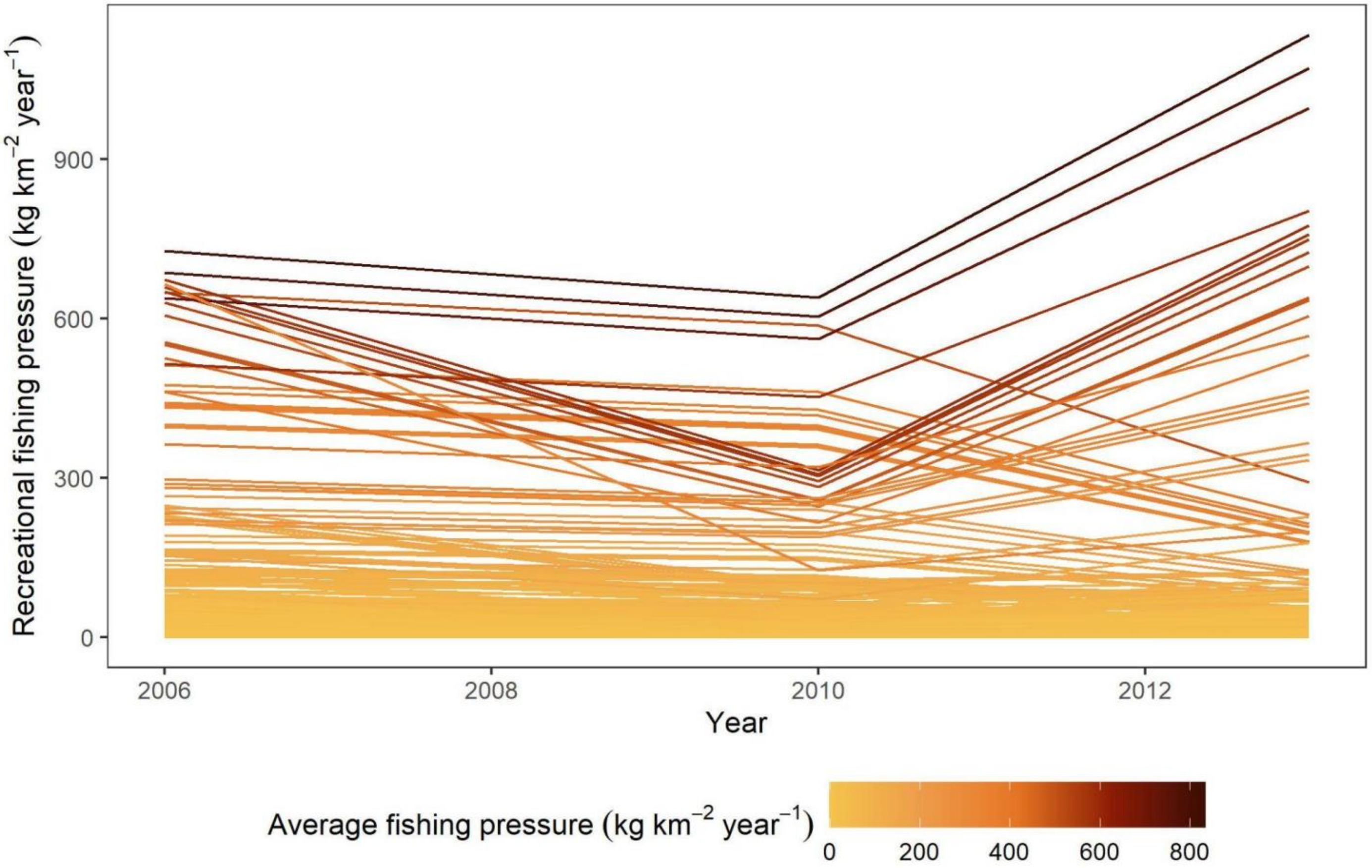


**Supplementary Fig. 23. Consistency in spatial pattern of recreational fishing pressure on perch**. Each line represents one ICES square, and shows estimated recreational fishing pressure per year for the three years for which data from questionnaires were available (2006, 2010 and 2013). Lines are coloured according to the average fishing pressure calculated across years for each location. In general, spatial patterns are broadly consistent over time and in line with average values. Note difference in scale compared with Supplementary Fig. 8 – fishing pressure from recreational fishing is much larger than that from commercial fisheries.

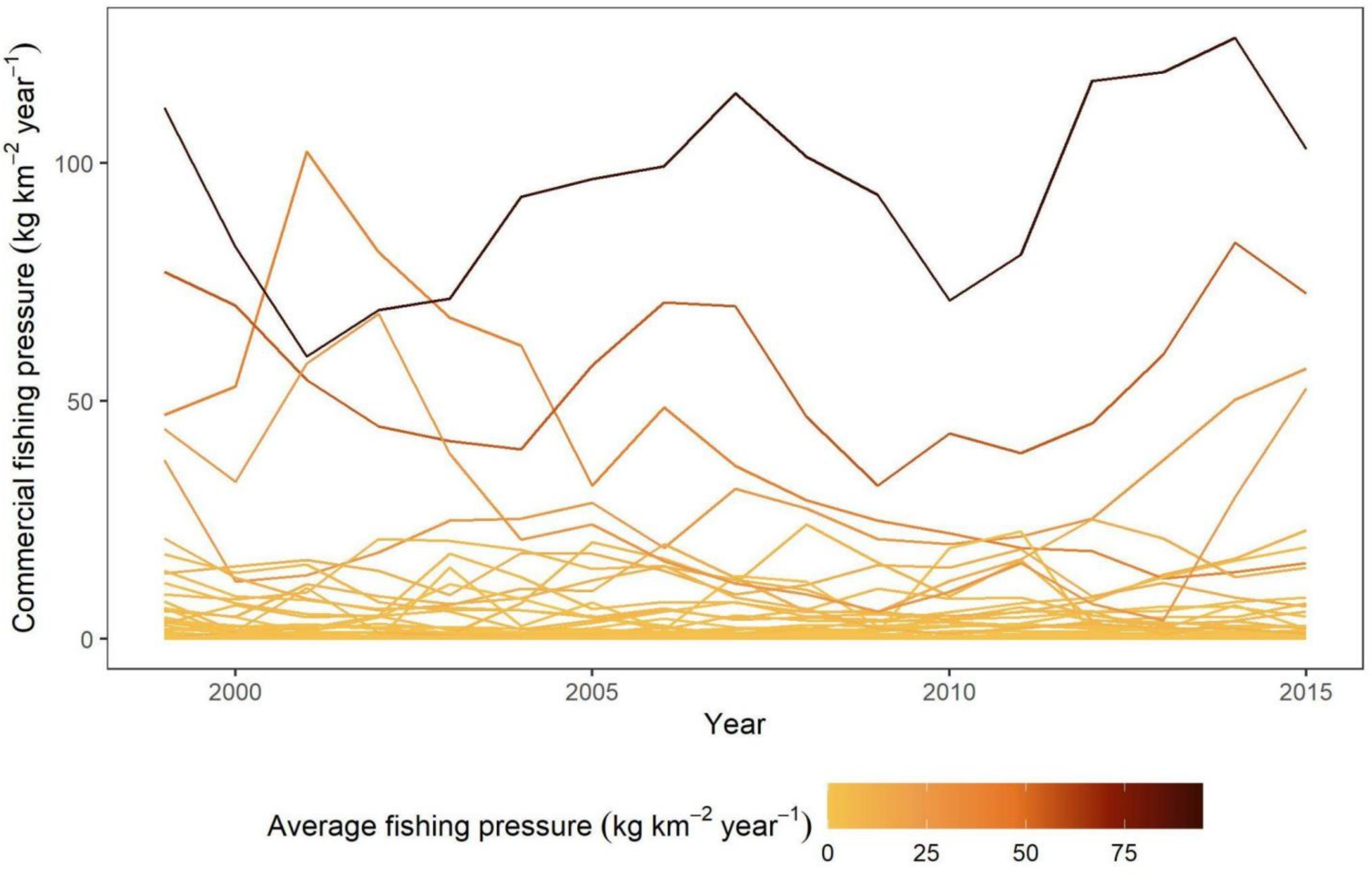


**Supplementary Fig. 24. Consistency in spatial pattern of commercial fishing pressure on perch.** Each line represents one ICES square, and shows estimated commercial fishing pressure per year from 1999 to 2015. Lines are coloured according to the average fishing pressure calculated across years for each location. In general, spatial patterns are broadly consistent over time and in line with average values. Note difference in scale compared with Supplementary Fig. 7 – fishing pressure from commercial fishing is much smaller than that from recreational fisheries.

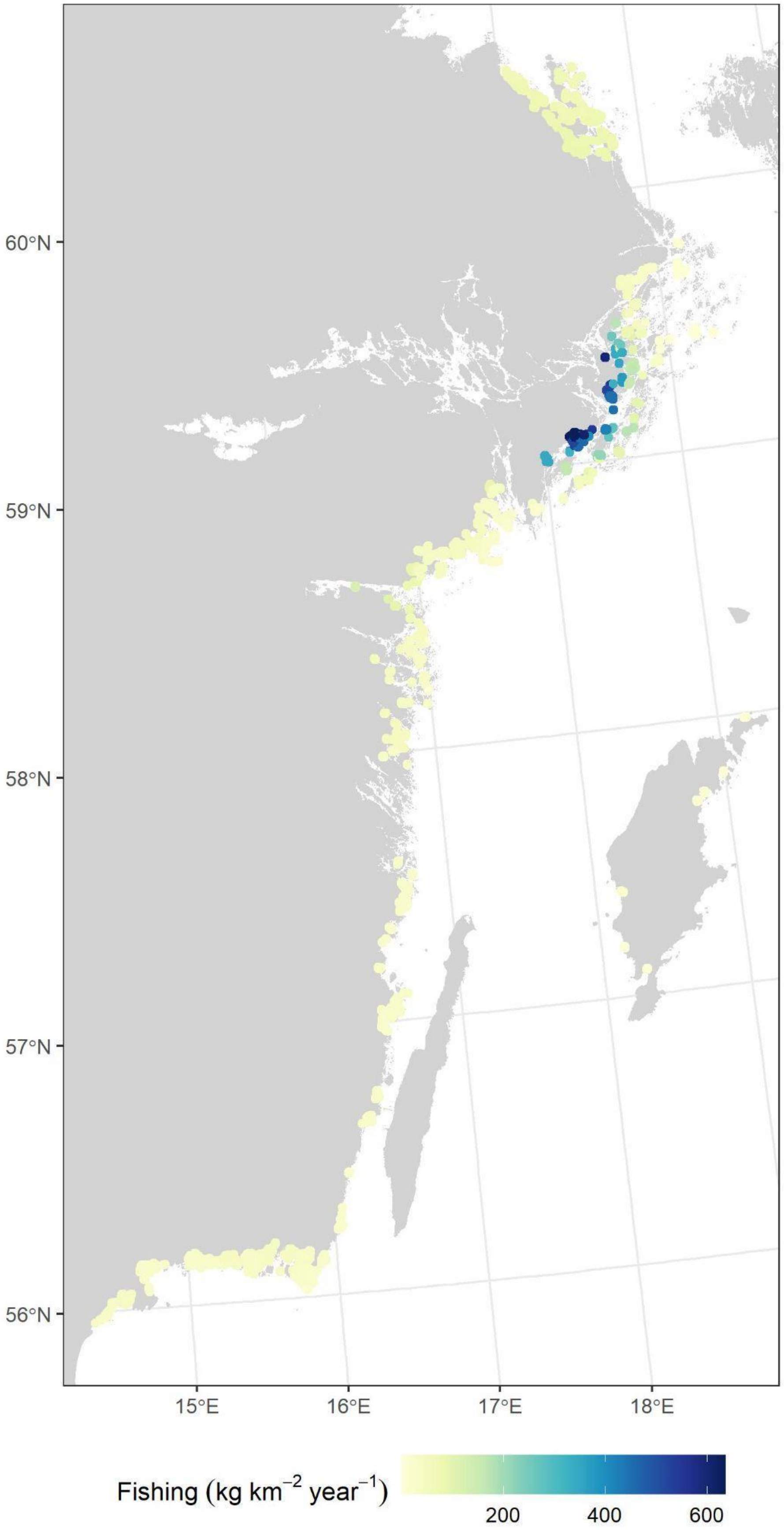


**Supplementary Fig. 25. Map of extracted values of fishing pressure for each detonation point.**

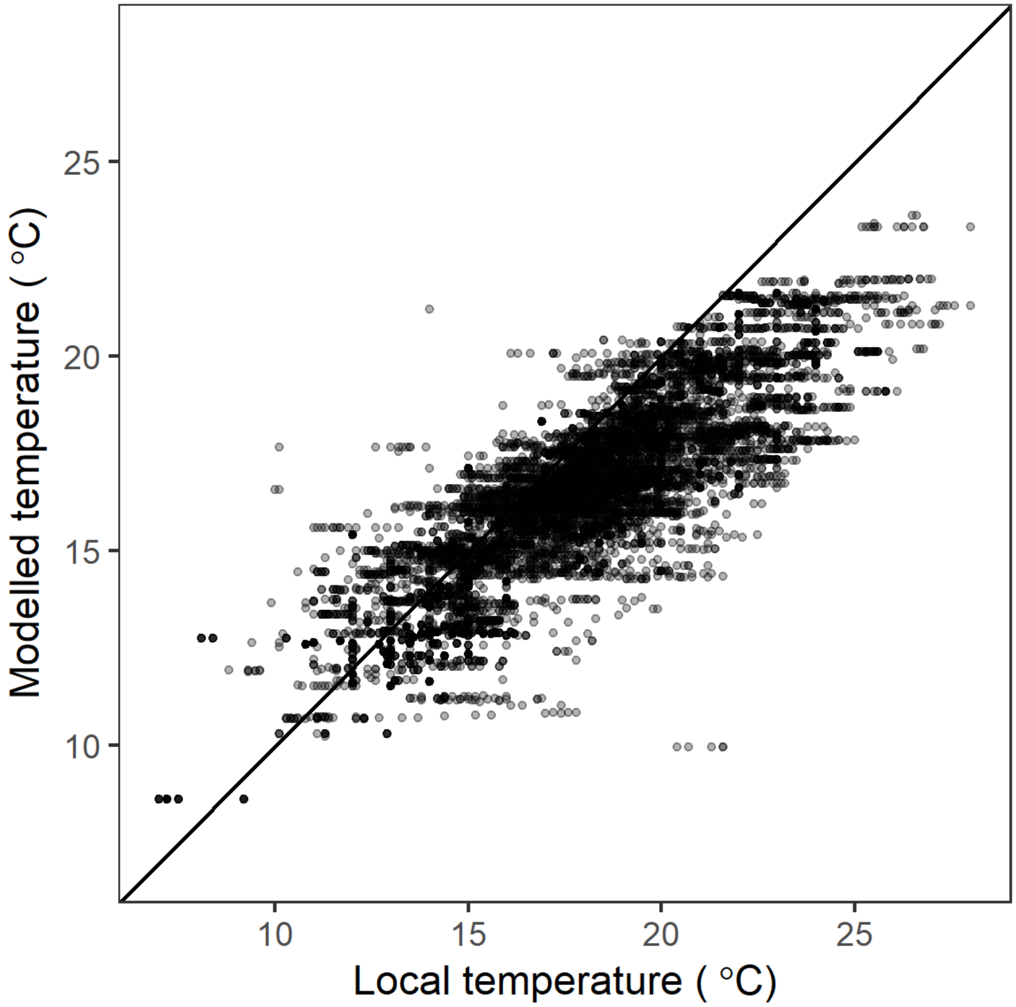


**Supplementary Fig. 26. Comparison of local temperatures and temperatures from satellite dataset.** Local temperatures are surface temperatures at the time of the detonations and modelled temperatures are extracted for the same day and location from the Copernicus Baltic Sea L4 dataset (https://doi.org/10.48670/moi-00156). There is a strong positive correlation between the two datasets ($r = 0.84$, $p < 0.001$). The plot suggests that the highest temperatures are slightly underestimated, likely because the resolution of the Copernicus dataset (ca. 1–2 km) is not sufficient for resolving small bays which may heat up more rapidly in summer.

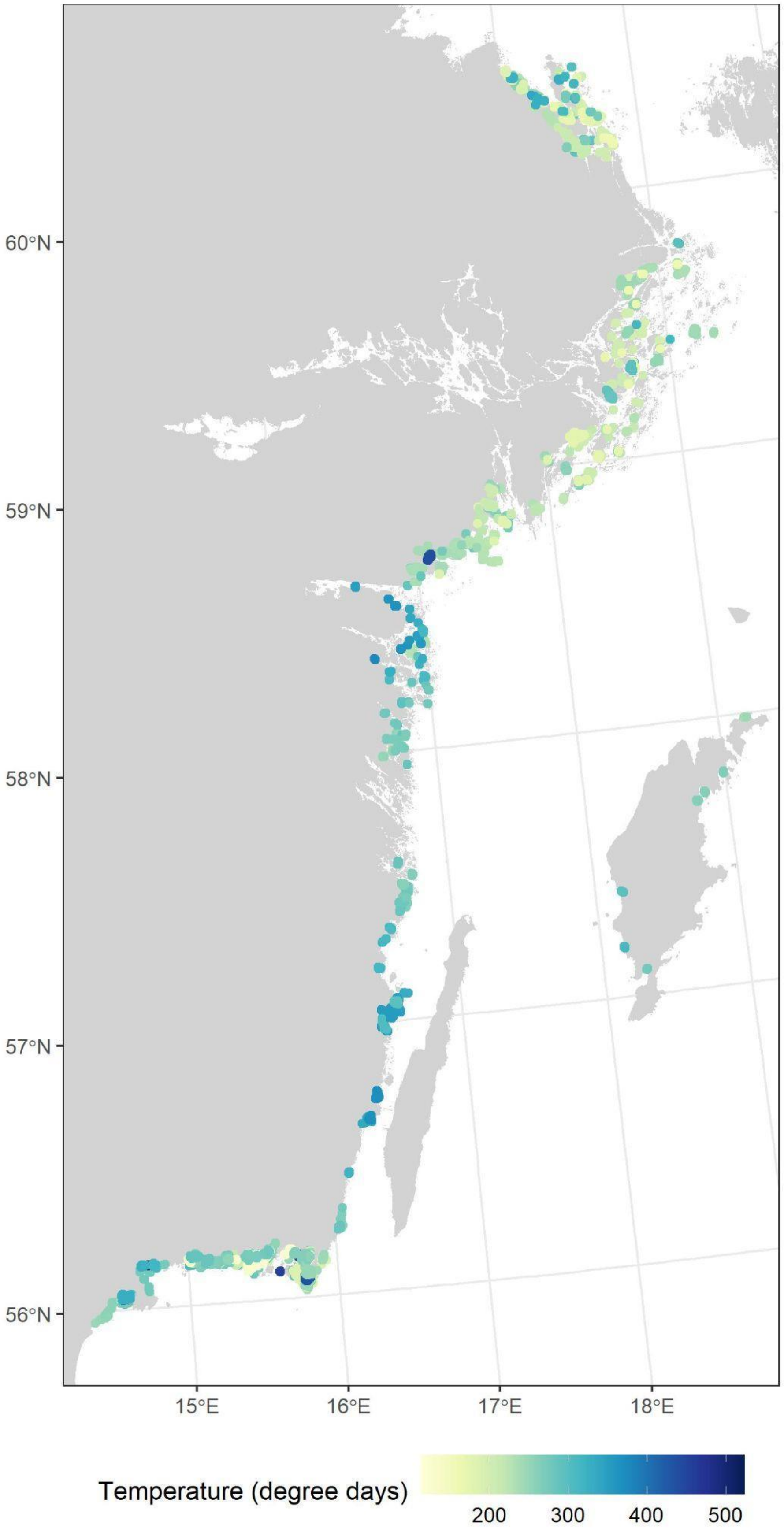


**Supplementary Fig. 27. Map of extracted values of temperature (degree days) for each detonation point.** Note that this varies between years, which is why close-by locations can have quite different values.

**Supplementary Table 8. Comparison of models with different random effect structures.** Comparison of models of predatory fish dominance (N = 3491), predatory fish densities (N = 7415), and stickleback densities (N = 7167) as a function of incoming stickleback (main effects of open sea stickleback densities, distance from the open sea, $\log_{10}$-transformed wave exposure, and an interaction between open sea stickleback densities and distance from the open sea) with different random effect structures. The comparison is based on Akaike's Information Criterion (AIC; lower relative to reference model better) and results from cross-validation: expected log predictive density (ELPD; higher is better). The cross-validation is based on 10 random folds that are identical across model structures. Unfortunately, ELPD could not be calculated for the most complex model with available computer resources due to memory allocation issues. Note that the AIC-values do not account for uncertainty in random effects. All models are fitted to the same dataset.

| Model | predatory fish dominance | | predatory fish density | | Stickleback density | |
|---|---|---|---|---|---|---|
| | ΔAIC | ELPD | ΔAIC | ELPD | ΔAIC | ELPD |
| Base model | 0 | -0.62 | 0 | -0.67 | 0 | -0.59 |
| Across-area random effect of year | -222.7 | -0.57 | -284.6 | -0.65 | -224.4 | -0.58 |
| Constant spatial random fields | -1195.7 | -0.37 | -977.3 | -0.62 | -1359.4 | -0.51 |
| Constant spatial random fields and across-area random effect of year | -1409.6 | -0.33 | -1234.7 | -0.61 | -1505.6 | -0.50 |
| Year-specific spatial random fields | -1427.1 | - | -1523.1 | - | -1854.5 | - |

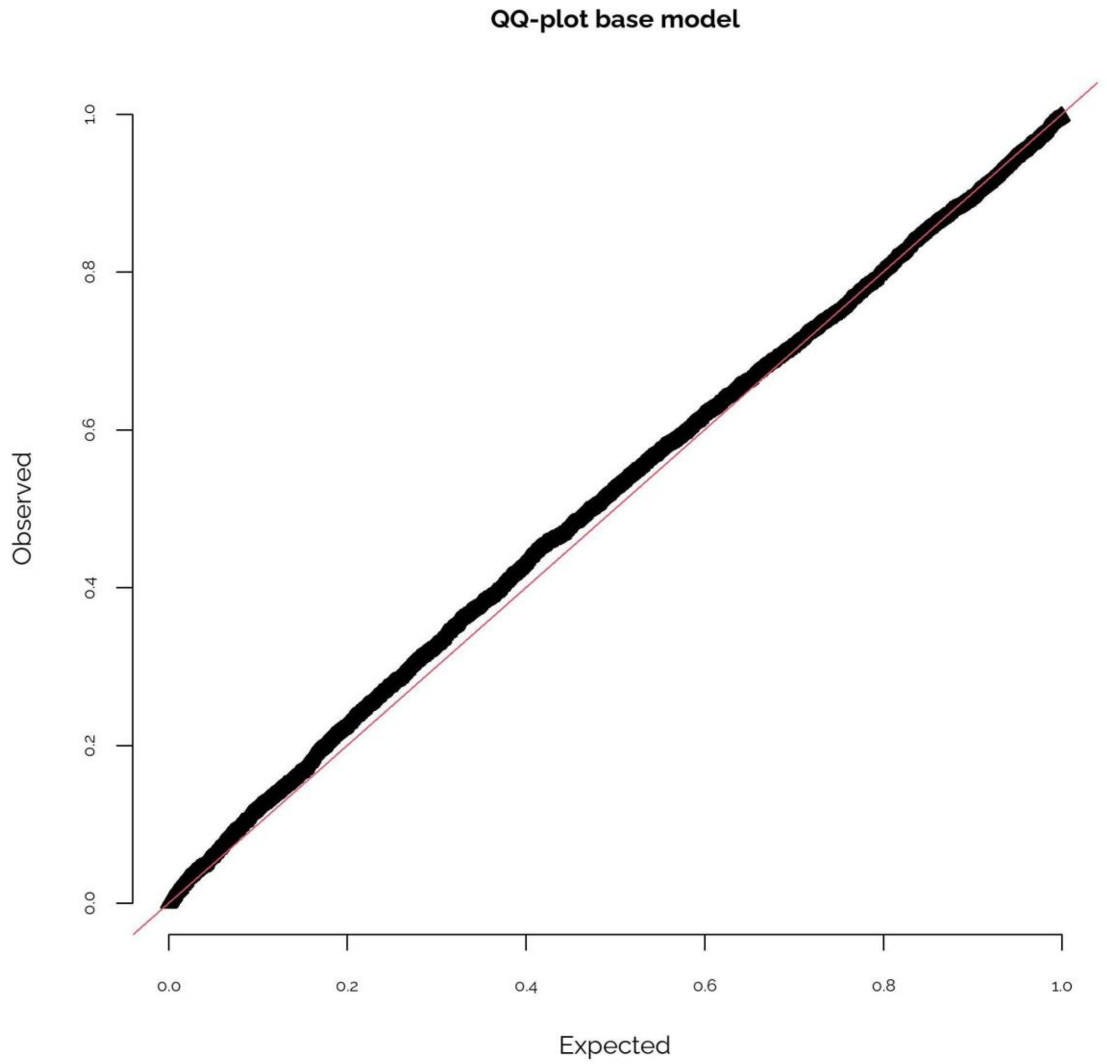


**Supplementary Fig. 28. QQ-plots for the full fitted model of relative predatory fish dominance, including main effects of open sea stickleback densities, distance from the open sea, $log_{10}$-transformed wave exposure, and an interaction between open sea stickleback densities and distance from the open sea, with year-specific spatial random fields.** The figure was produced using the R-package DHARMa (Hartig, 2022). The tests provided by DHARMa pointed to some deviations from the expected error distributions, but these deviations were small, as shown in the QQ-plot. While most of the plots of predictors against residuals showed signs of a pattern, this pattern was often wobbly but did not point to any alternative functional forms (such as a quadratic relationship). Variance inflation factors were below 2 for all variables, and there was thus no indication of multicollinearity. Finally, as could be expected, the models that did not include random spatial fields showed clear signs of spatial autocorrelation in the residuals (Moran's I = 0.21/0.17 [no random effect of year/random effect of year], $p < 0.001$). After refitting the models with random spatial fields, there was no spatial autocorrelation in the residuals (Moran's I = 0.01/0.01/0.01 p = 0.19/0.21/0.19 [spatial random fields only/spatial random fields + random year effect/spatio-temporal random fields]).

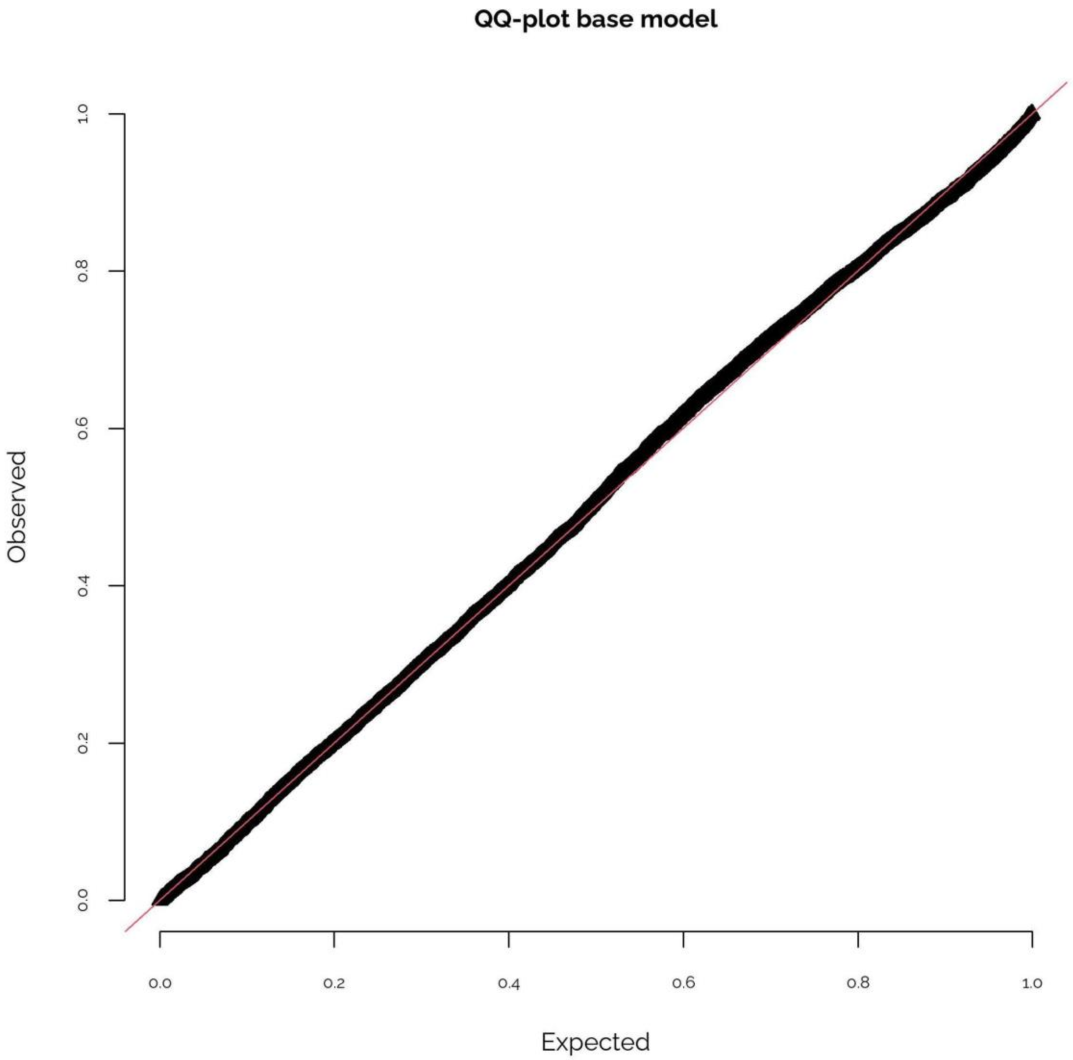


**Supplementary Fig. 29. QQ-plot for the full fitted model of predatory fish densities, including main effects of open sea stickleback densities, distance from the open sea, $log_{10}$-transformed wave exposure, and an interaction between open sea stickleback densities and distance from the open sea, with year-specific spatial random fields.** The figure was produced using the R-package DHARMa (Hartig, 2022). The tests provided by DHARMa pointed to some deviations from the expected error distributions in some of the model representations (i.e. the different random effect structures), but these deviations were small, as shown in the QQ-plot. While most of the plots of predictors against residuals showed signs of a pattern, this pattern did not point to any alternative functional forms (such as a quadratic relationship). Variance inflation factors were below 2 for all variables, and there was thus no indication of multicollinearity. Finally, as could be expected, the models that did not include random spatial fields showed signs of spatial autocorrelation in the residuals (Moran's I = 0.13/0.11 [no random effect of year/random effect of year], $p < 0.001$). After refitting the models using sdmTMB, spatial autocorrelation remained but was reduced (Moran's I = 0.06/0.05/0.05 $p < 0.001$ [spatial random fields only/spatial random fields + random year effect/spatio-temporal random fields]). Correctly specified spatial models may still show spatial autocorrelation in the residuals, but will still provide more accurate standard errors (Beale et al., 2010).

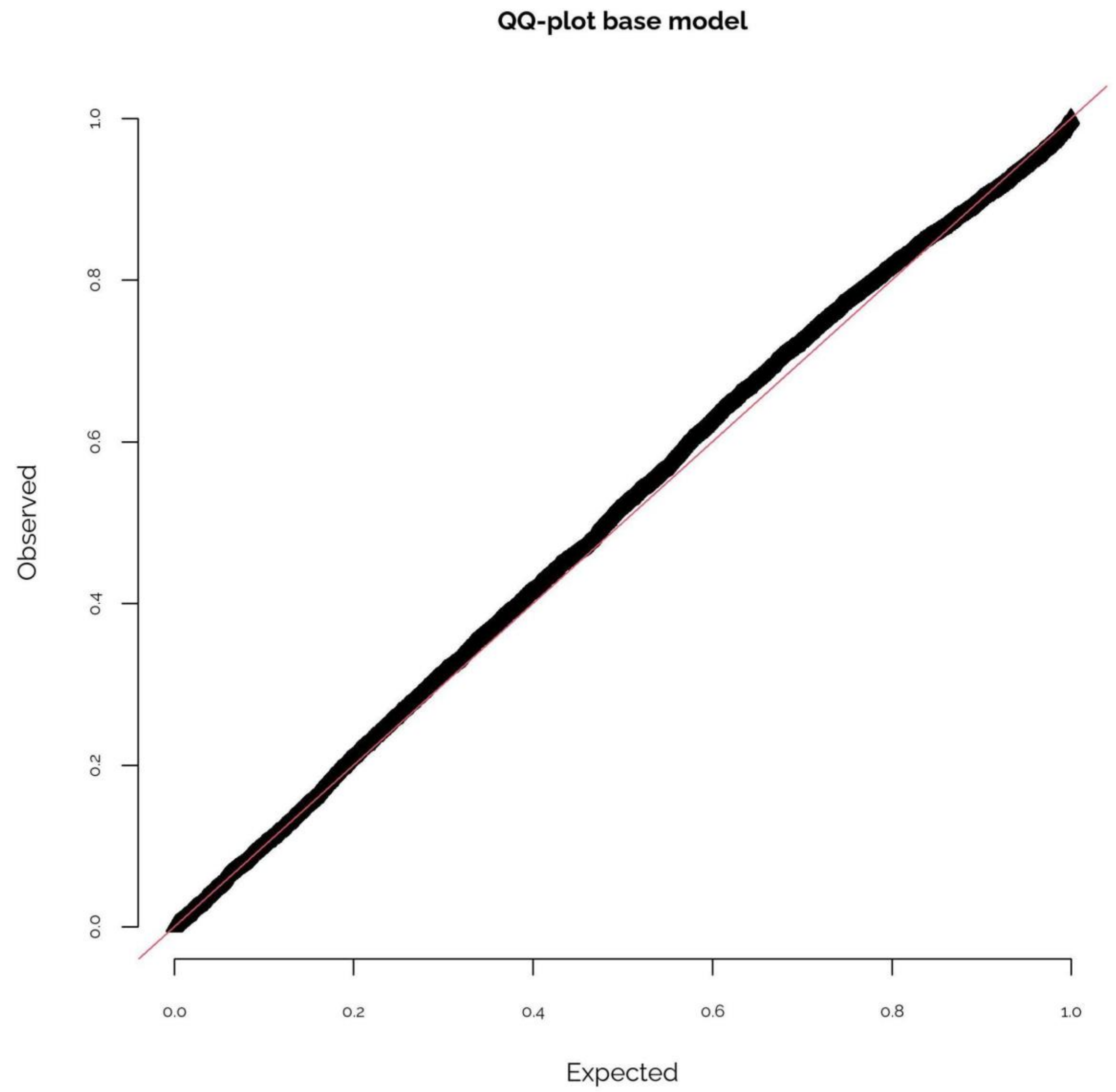


**Supplementary Fig. 30. QQ-plots for the full fitted model of stickleback densities, including main effects of open sea stickleback densities, distance from the open sea, $\log_{10}$-transformed wave exposure, and an interaction between open sea stickleback densities and distance from the open sea, with year-specific spatial random fields.** The figure was produced using the R-package DHARMa (Hartig, 2022). The tests provided by DHARMa pointed to some deviations from the expected error distributions in some of the model representations (i.e. the different random effect structures), but these deviations were small, as shown in the QQ-plot. While most of the plots of predictors against residuals showed signs of a pattern, this pattern did not point to any alternative functional forms (such as a quadratic relationship). Variance inflation factors were below 2 for all variables, and there was thus no indication of multicollinearity. Finally, as could be expected, the models that did not include random spatial fields showed signs of spatial autocorrelation in the residuals (Moran's I = 0.14/0.12 [no random effect of year/random effect of year], $p < 0.001$). After refitting the models using sdmTMB, spatial autocorrelation remained but was reduced (Moran's I = 0.04/0.03/0.04 $p < 0.001$ [spatial random fields only/spatial random fields + random year effect/spatio-temporal random fields]). Correctly specified spatial models may still show spatial autocorrelation in the residuals, but will still provide more accurate standard errors (Beale et al., 2010).

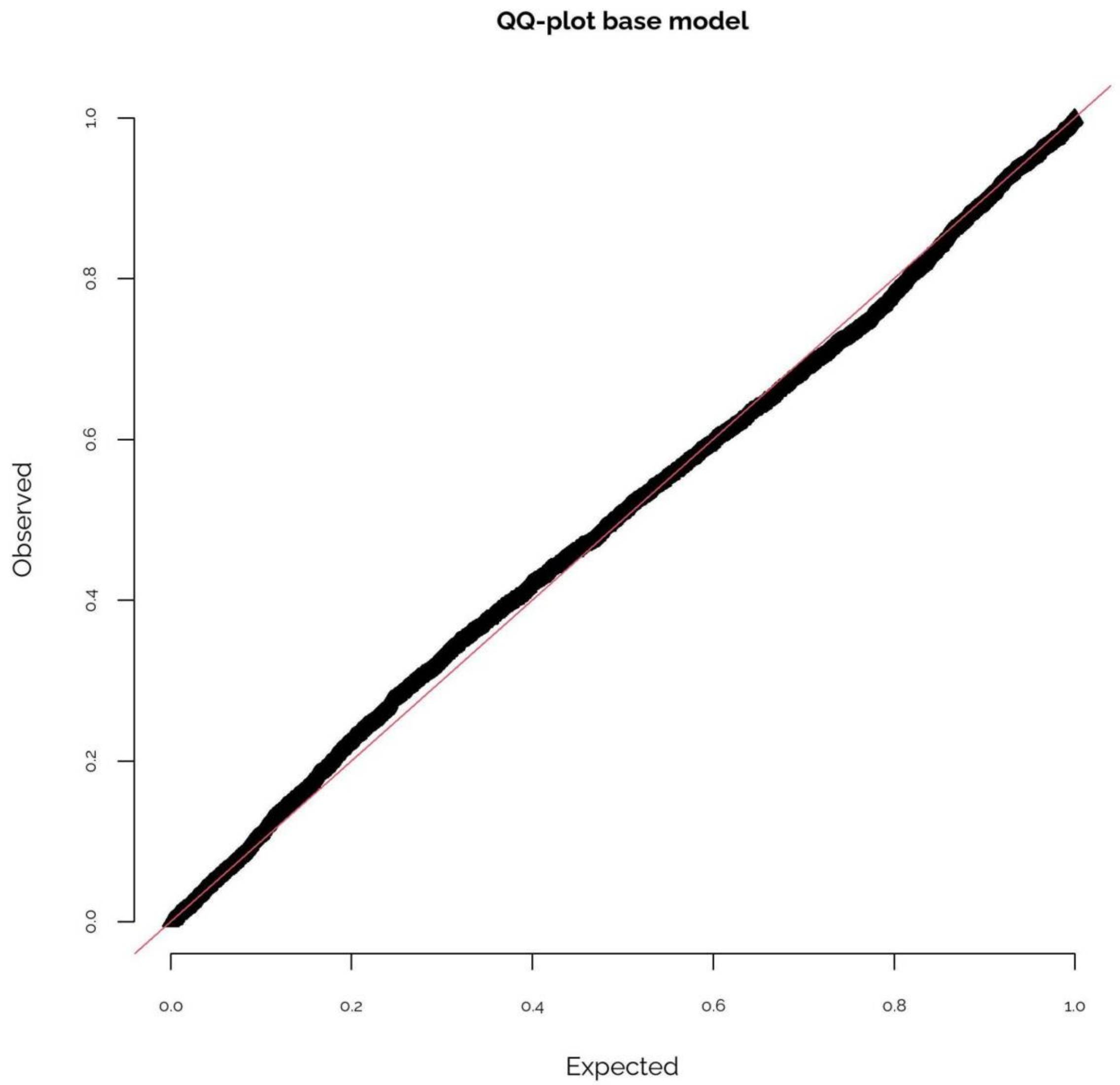


**Supplementary Fig. 31. QQ-plots for the full fitted model of predatory fish dominance, including main effects of open sea stickleback densities, distance from the open sea, $log_{10}$-transformed wave exposure, an interaction between open sea stickleback densities and distance from the open sea, predation, fishing, temperature, connectivity (network representation, 3.5 cut-off since this was the best representation for predatory fish dominance), interactions between connectivity and predation, between connectivity and fishing, and between temperature and distance from the open sea, as well as a random effect of year.** The figure was produced using the R-package DHARMa (Hartig, 2022). The tests provided by DHARMa pointed to some deviations from the expected error distributions in some of the model representations, but these deviations were small, as shown in the QQ-plot. While most of the plots of predictors against residuals showed signs of a pattern, this pattern did not point to any alternative functional forms (such as a quadratic relationship). Variance inflation factors were below 3 for all variables (majority below 2), except for fishing where they were 2.6–5.5 depending on the connectivity representation (see Supplementary Tables 9–12 for between-variable correlations). We thus re-ran the models not including fishing effects to confirm that our conclusions were still supported, which they were. Finally, there were signs of spatial autocorrelation in the residuals (Moran's I = 0.17–0.18, $p < 0.001$).

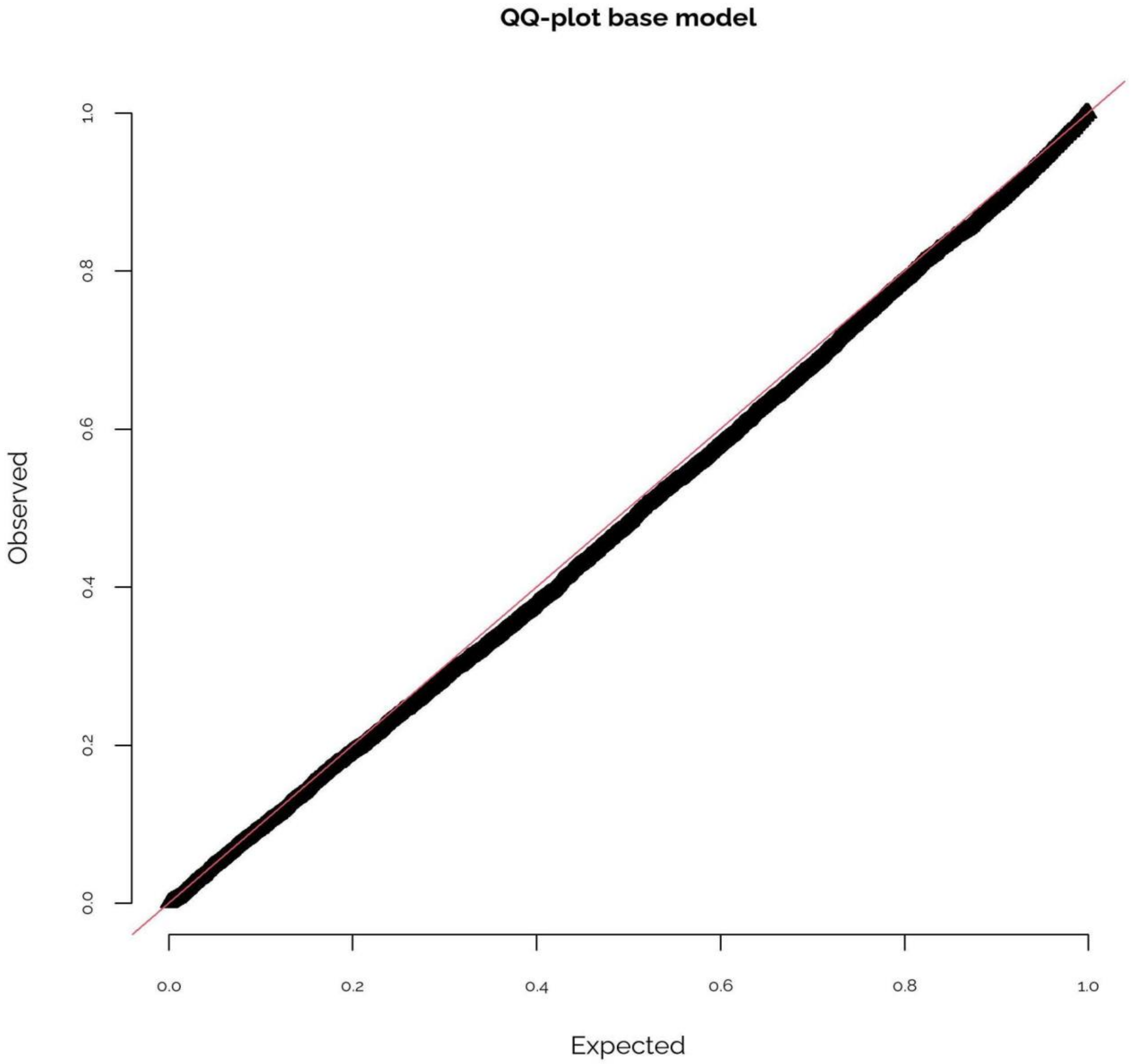


**Supplementary Fig. 32. QQ-plots for the full fitted model of predatory fish densities, including main effects of open sea stickleback densities, distance from the open sea, $log_{10}$-transformed wave exposure, an interaction between open sea stickleback densities and distance from the open sea, predation, fishing, temperature, connectivity (weighted sum of all available habitat within a 10 km radius and using a 3.2 cut-off for wave exposure since this was the best representation for predatory fish densities), interactions between connectivity and predation, connectivity and fishing, and temperature and distance from the open sea, as well as a random effect of year.** The figure was produced using the R-package DHARMa (Hartig, 2022). The tests provided by DHARMa pointed to some deviations from the expected error distributions in some of the model representations, but these deviations were small, as shown in the QQ-plot. While most of the plots of predictors against residuals showed signs of a pattern, this pattern did not point to any alternative functional forms (such as a quadratic relationship). Variance inflation factors were below 3 for all variables (majority below 2), except for fishing where they were 1.6–6.6 depending on the connectivity representation (see Supplementary Tables 9–12 for between-variable correlations). We thus re-ran the models not including fishing effects to confirm that our conclusions were still supported, which they were. Finally, there were signs of spatial autocorrelation in the residuals (Moran's I = 0.11, $p < 0.001$).

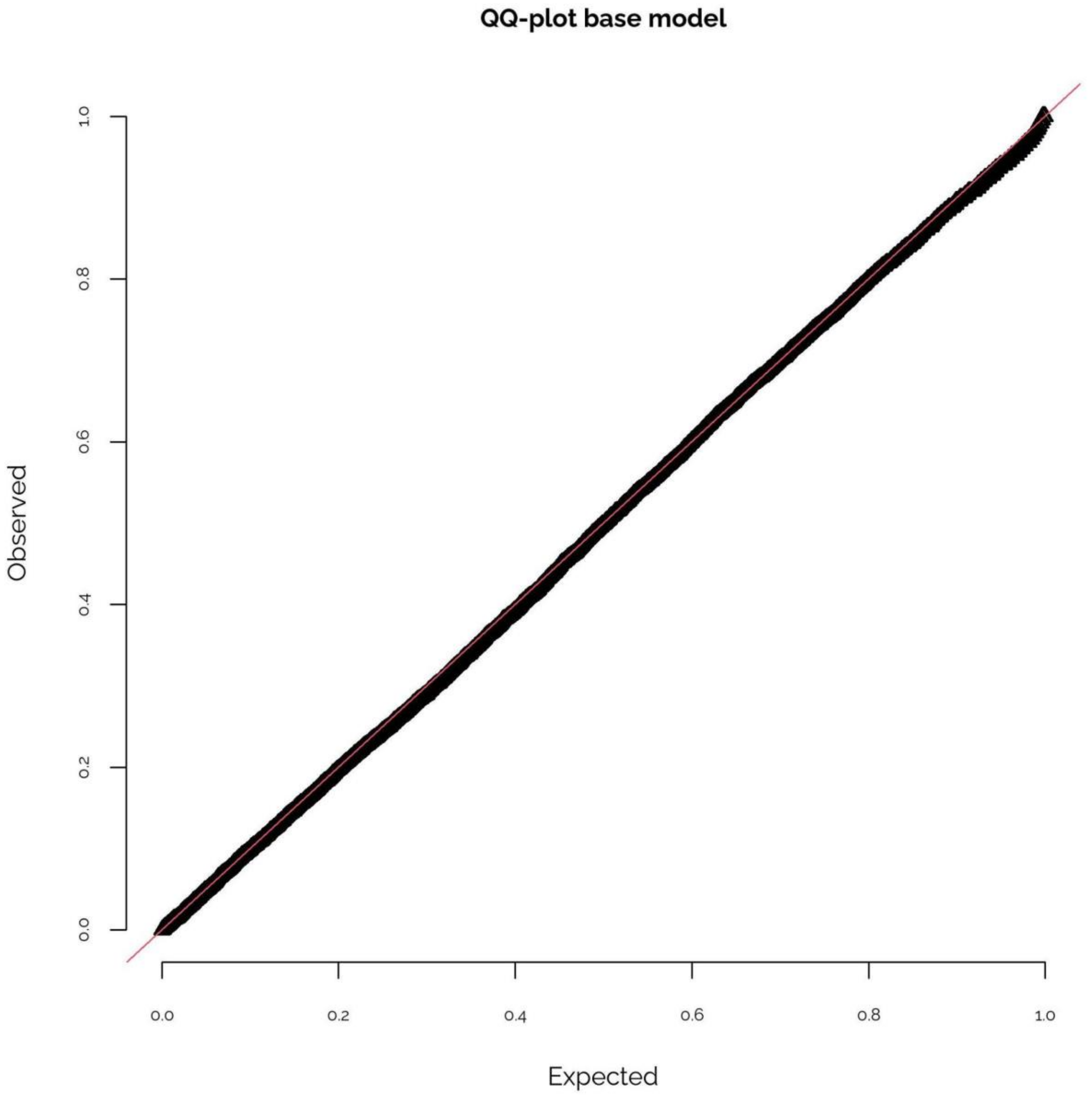


**Supplementary Fig. 33. QQ-plots for the full fitted model of stickleback densities, including main effects of open sea stickleback densities, distance from the open sea, $log_{10}$-transformed wave exposure, an interaction between open sea stickleback densities and distance from the open sea, predation, fishing, temperature, connectivity (weighted sum of all available habitat within a 10 km radius and using a 3.5 cut-off for wave exposure since this was the best representation for stickleback densities), interactions between connectivity and predation, connectivity and fishing, and temperature and distance from the open sea, as well as a random effect of year.** The figure was produced using the R-package DHARMa (Hartig, 2022). The tests provided by DHARMa pointed to some deviations from the expected error distributions in some of the model representations, but these deviations were small, as shown in the QQ-plot. While most of the plots of predictors against residuals showed signs of a pattern, this pattern did not point to any alternative functional forms (such as a quadratic relationship). Variance inflation factors were below 3 for all variables (majority below 2), except for fishing where they were 1.6–5.6 depending on the connectivity representation (see Supplementary Tables 9–12 for between-variable correlations). We thus re-ran the models not including fishing effects to confirm that our conclusions were still supported, which they were. Finally, there were signs of spatial autocorrelation in the residuals (Moran's I = 0.12–0.13, $p < 0.001$).

**Supplementary Table 9. Correlations between explanatory variables (N = 7415).** Correlations with $p < 0.05$ are indicated in bold. Here connectivity is the weighted sum of all available habitat within a 10 km radius and using a 3.5 cut-off for $\log_{10}$-transformed wave exposure. stickleback = offshore densities of mature stickleback, WE = $\log_{10}$-transformed wave exposure.

| | connectivity | stickleback | distance | WE | predation | fishing |
|---|---|---|---|---|---|---|
| stickleback | **0.08** | | | | | |
| distance | **0.41** | **0.14** | | | | |
| WE | **-0.26** | **0.04** | **-0.24** | | | |
| predation | **0.10** | **0.38** | **0.19** | **-0.07** | | |
| fishing | **-0.03** | **0.19** | **0.35** | **-0.09** | -0.01 | |
| temperature | **0.03** | **-0.19** | **-0.23** | -0.02 | **0.18** | **-0.19** |

**Supplementary Table 10. Correlations between explanatory variables (N = 7415).** Correlations with $p < 0.05$ are indicated in bold. Here connectivity is the weighted sum of all available habitat within a 10 km radius and using a 3.2 cut-off for $\log_{10}$-transformed wave exposure. stickleback = offshore densities of mature stickleback, WE = $\log_{10}$-transformed wave exposure.

| | connectivity | stickleback | distance | WE | predation | fishing |
|---|---|---|---|---|---|---|
| stickleback | **0.07** | | | | | |
| distance | **0.12** | **0.14** | | | | |
| WE | **-0.23** | **0.04** | **-0.24** | | | |
| predation | **0.08** | **0.38** | **0.19** | **-0.07** | | |
| fishing | **-0.09** | **0.19** | **0.35** | **-0.09** | -0.01 | |
| temperature | **0.16** | **-0.19** | **-0.23** | -0.02 | **0.18** | **-0.19** |

**Supplementary Table 11. Correlations between explanatory variables (N = 7415).** Correlations with $p < 0.05$ are indicated in bold. Here connectivity is connected habitat based on the network approach with a 3.5 cut-off for $\log_{10}$-transformed wave exposure. stickleback = offshore densities of mature stickleback, WE = $\log_{10}$-transformed wave exposure.

| | connectivity | stickleback | distance | WE | predation | fishing |
|---|---|---|---|---|---|---|
| stickleback | -0.02 | | | | | |
| distance | **0.17** | **0.14** | | | | |
| WE | **-0.21** | **0.04** | **-0.24** | | | |
| predation | **0.09** | **0.38** | **0.19** | **-0.07** | | |
| fishing | **-0.17** | **0.19** | **0.35** | **-0.09** | -0.01 | |
| temperature | **0.10** | **-0.19** | **-0.23** | -0.02 | **0.18** | **-0.19** |

**Supplementary Table 12**. **Correlations between explanatory variables (N = 7415).** Correlations with $p < 0.05$ are indicated in bold. Here connectivity is connected habitat based on the network approach with a 3.2 cut-off for $log_{10}$-transformed wave exposure. stickleback = offshore densities of mature stickleback, WE = $log_{10}$-transformed wave exposure.

| | connectivity | stickleback | distance | WE | predation | fishing |
|---|---|---|---|---|---|---|
| stickleback | 0.00 | | | | | |
| distance | **-0.10** | **0.14** | | | | |
| WE | **-0.17** | **0.04** | **-0.24** | | | |
| predation | **0.09** | **0.38** | **0.19** | **-0.07** | | |
| fishing | **-0.16** | **0.19** | **0.35** | **-0.09** | -0.01 | |
| temperature | **0.23** | **-0.19** | **-0.23** | -0.02 | **0.18** | **-0.19** |
